# Optogenetic cochlear stimulation evokes midbrain activity with near-physiological temporal fidelity

**DOI:** 10.64898/2026.05.16.724905

**Authors:** Elisabeth Koert, Jonathan Götz, Niels Albrecht, Anna Vavakou, Bettina Julia Wolf, Tobias Moser

## Abstract

When hearing fails, stimulation of the auditory nerve by electrical cochlear implants (eCIs) partially restores hearing, with most eCI users achieving open speech understanding. However, the broad current spread from each electrode limits frequency coding and speech understanding in daily situations with background noise. Spatially confined optogenetic stimulation by future optical cochlear implants (oCIs) improves frequency coding but millisecond closing kinetics of channelrhodopsins (ChRs) might limit temporal coding. Here, we evaluated the utility of fast-closing f-Chrimson for processing temporal information in the auditory system of Mongolian gerbils. We recorded neural activity in the inferior colliculus evoked by f-Chrimson-mediated optogenetic stimulation of the cochlea. F-Chrimson enabled energy-efficient stimulation of the auditory pathway at rates ≥150 Hz, outperforming the slower ChR variants CatCh (blue) and ChReef (green). Energy thresholds for activation of the auditory pathway were in the low µJ range, between ChReef (sub-µJ) and CatCh. Dynamic range and frequency selectivity were comparable to previous observations with CatCh and outperformed electrical stimulation. In conclusion, employing fast-gating ChRs harnesses improved spectral coding without degrading temporal coding.

**The Paper Explained:** 

**Problem:** Electrical cochlear implants (eCIs) partially restore speech comprehension in most of 1 million otherwise severely deaf people. However, most CI-users face challenges hearing in daily situations. Spectrally more selective stimulation of the auditory nerve by optical cochlear implants (oCIs) promises to overcome this limitation. However, the closing kinetics of channelrhodopsins (ChR) limit the temporal bandwidth of bionic sound coding. Improving the ChR properties and evaluating temporal coding remain major objectives for developing hearing restoration by oCI.

**Results:** Here, we evaluate the utility of waveguide-based oCI using the fast-closing ChR Chrimson (f-Chrimson) for encoding of temporal, spectral and intensity information by multi-electrode-array (MEA) recordings from the midbrain. We compare f-Chrimson-mediated bionic coding to acoustic coding as well as to previous data acquired with optogenetic stimulation using other ChRs and with electrical stimulation. F-Chrimson enabled energy-efficient stimulation of the auditory pathway at rates ≥150 Hz, outperforming the slower ChR variants CatCh (blue) and ChReef (green). Intensity and frequency coding were comparable to previous observations with CatCh and outperformed electrical stimulation.

**Impact:** This study demonstrates near physiological temporal coding with the fast-closing ChR f-Chrimson, indicating that improved spectral coding by oCI is not traded off by poor temporal fidelity.

## Introduction

Hearing loss is the most common sensory disorder, with ∼0.5 billion people requiring intervention (disabling hearing loss, (WHO, 2024)). Children require hearing to acquire vocal speech. Unaided hearing loss in adults limits professional and private activities, frequently causes social isolation and poses a risk for depression and cognitive decline (WHO, 2024). Until recently, no causal therapies were available for the most frequent form, sensorineural hearing loss resulting from malfunction of the cochlea and/or the auditory nerve. Now, with promising results of several clinical trials on gene replacement therapy for *OTOF*-related auditory synaptopathy (e.g. Lv *et al*, 2024; Qi *et al*, 2024; Valayannopoulos *et al*, 2025) there is hope for a change at least for genetic forms of deafness (Fan *et al*, 2025; Moser *et al*, 2024). Yet, even there, with more than 150 deafness genes, it seems not practically feasible to provide tailored therapies for many deafness genes. This is primarily due to early degeneration of the sensory cells, demonstrated by mouse models, precluding gene therapy or at least a postnatal window of opportunity. As a result, we project that the number of genes for which gene therapy will become available will probably not exceed a dozen (Vona *et al*, 2025). Considering this and given the many other indications for CI implantations, one-suits all approaches such as hearing aids and CIs will remain viable treatment options for the coming decades, while alternative solutions including regenerative medicine are being developed (Wrobel *et al*, 2021).

Yet, there remains a major unmet medical need to improve hearing restoration with these devices and the cochlear implant (CI) in particular (Wolf *et al*, 2022). CI-patients demand better functional outcomes in real-world settings such as communication in challenging acoustic environments in daily life (Hunniford *et al*, 2023). Many of the current challenges are attributed to the broad spread of current from each electrode contact in the cochlear saline. This activates large numbers of the tonotopically organized spiral ganglion neurons (SGNs), which results in poor coding frequency and intensity information (Shannon, 1983; Kral *et al*, 1998; Miller *et al*, 2006). As light can be spatially confined, optical stimulation could improve frequency coding and, consequently, hearing restoration for CI users (Izzo *et al*, 2006; Hernandez *et al*, 2014). While direct optical stimulation of SGNs (Izzo *et al*, 2006) remains controversial (Azimzadeh *et al*, 2025), optogenetic stimulation of SGNs has been demonstrated in several species and for various ChRs (Hernandez *et al*, 2014; Wrobel *et al*, 2018; Duarte *et al*, 2018; Thompson *et al*, 2020; Alekseev *et al*, 2025). Indeed, blue and green-light optogenetic SGN stimulation showed improved spectral selectivity compared to eCI stimulation in previous studies (Dieter *et al*, 2019, 2020; Keppeler *et al*, 2020; Alekseev *et al*, 2025) as well as the companion (Albrecht et al.). These studies addressed the potential limitation of the temporal fidelity of optogenetic SGN stimulation resulting from the closing kinetics of ChRs (Mager *et al*, 2018; Keppeler *et al*, 2018; Bali *et al*, 2021; Mittring *et al*, 2023; Alekseev *et al*, 2025; Roos *et al*, 2026). Efforts have been undertaken to speed up ChR closing kinetics for optogenetic hearing restoration (Mager *et al*, 2018; Keppeler *et al*, 2018; Mittring *et al*, 2023; Roos *et al*, 2026). Fast versions of the orange-light activated ChR Chrimson (Klapoetke *et al*, 2014; Mager *et al*, 2018) have been characterized in mice at the level of auditory brainstem response and single SGN firing (Mager *et al*, 2018; Bali *et al*, 2021). Yet a study, that would investigate the utility of fast (f)-Chrimson for time, intensity and frequency coding in a more translational animal model remained to be performed.

Here, we employed the Mongolian gerbil, with a cochlear size and hearing frequency range closer to human for characterizing f-Chrimson mediated SGN stimulation by recordings of auditory brainstem responses and multiunit (MU) activity in the central nucleus of the inferior colliculus (ICC). The inferior colliculus provides access to all three attributes of the auditory code (Drotos & Roberts, 2024) and was used in comparing the optogenetic coding with data on acoustic and electrical coding obtained in previous work. We find that the temporal fidelity of f-Chrimson mediated SGN stimulation outperforms that obtained with the slower ChRs CatCh and ChReef while maintaining comparable frequency selectivity and dynamic range.

## Results

### Postnatal AAV mediated gene transfer of f-Chrimson into gerbil spiral ganglion neurons (SGNs) allows for activation of the auditory pathway

We injected AAV-PHP.B carrying f-Chrimson under the control of the human synapsin into the cochlea of Mongolian gerbils at P5-P9. In animals aged 8 weeks or older, we recorded auditory brainstem responses (ABRs) with acoustic and optical stimulation (Fig. 1). First, we delivered 0.1 ms click stimuli from a free-field speaker to the AAV-f-Chrimson injected ear to determine the acoustic auditory brainstem response (aABR) thresholds that amounted 32.5 ± 6 dB SPL and compares favourably to previous reports (Wrobel et al, 2018; Michael et al, 2023). We did not find significant differences in aABR threshold when separating the animals by gene therapy success (Fig. S1). We then performed a bullostomy and inserted a 200 µm optical fiber coupled to a 594 nm laser into the round window for recording ABRs to 1 ms optical stimulation (oABR). The surgical approach is illustrated in Fig. S2A.

**Figure 1.**
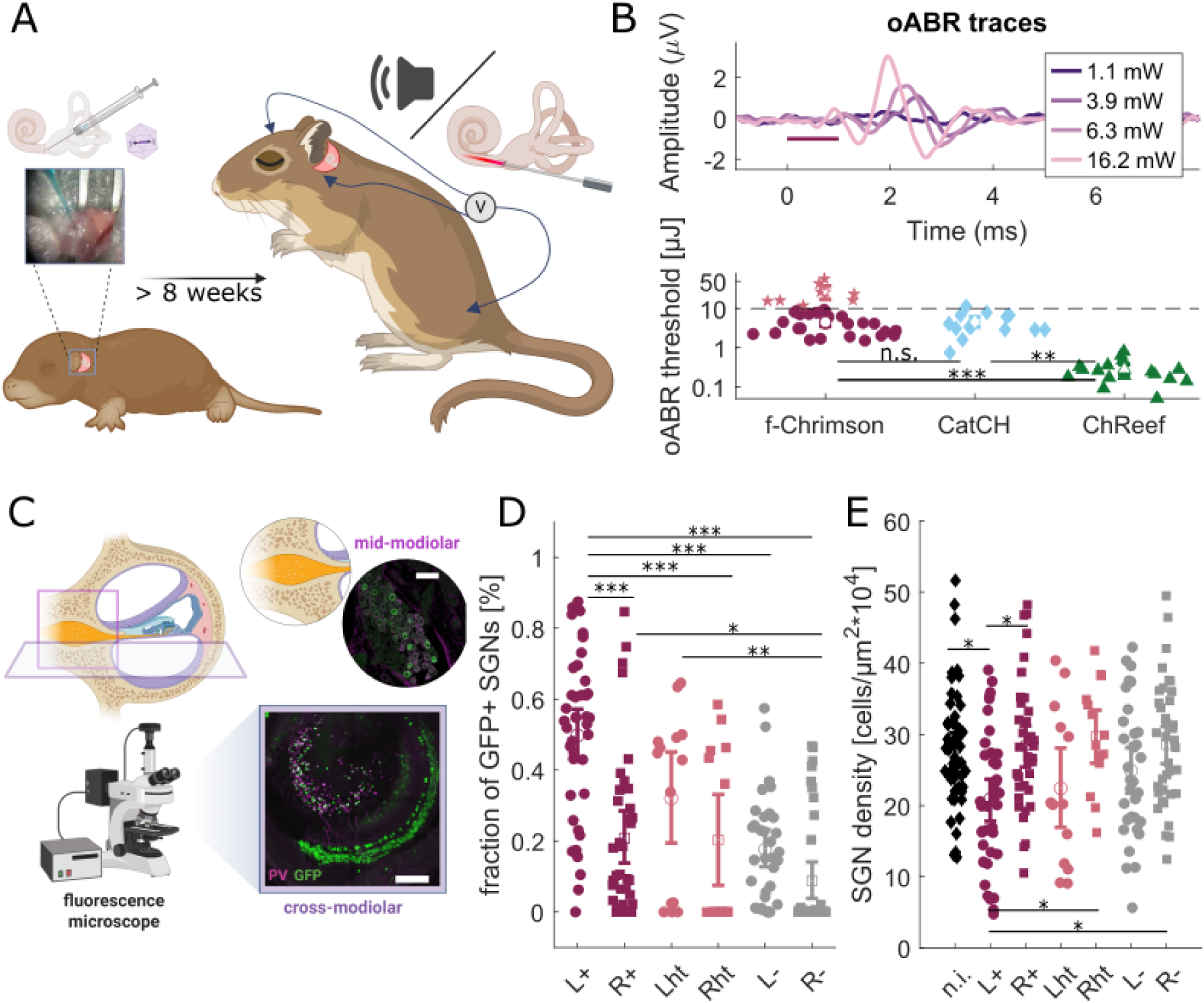
Postnatal AAV-mediated gene transfer of f-Chrimson into gerbil spiral ganglion neurons enables optogenetic activation of the auditory pathway. **(A)** Mongolian gerbils were postnatally injected with AAV-PHP.B carrying f-Chrimson under the control of the human synapsin promotor. At >8 weeks of age, auditory brainstem responses were recorded with acoustic (aABR) and optical (oABR) stimulation of the cochlea. (Created in BioRender. Koert, E. (2026) https://BioRender.com/22retyh) **(B)** Top: exemplary oABR response to 594 nm light pulses (1 ms, magenta bar) at the indicated radiant flux. Bottom: Thresholds for visual detection of the first wave. f-Chrimson animals with high thresholds (pink stars, >10 µJ) were excluded from the further experiments. For comparison, we included previously published thresholds for CatCh (Wrobel et al, 2018) and for ChReef from (Alekseev et al, 2025) and from the companion study (Albrecht et al.). **(C)** Cochleae were extracted, fixed, sliced (cross-modiolar vibratome sections or mid-modiolar cryostat sections) and immunolabeled for parvalbumin (PV) and GFP. (Created in BioRender. Koert, E. (2026) https://BioRender.com/eb414zj, scalebar: 50 µm for mid-modiolar, 200 µm for cross-modiolar) **(D, E)** Fraction of GFP^+^ SGNs and SGN density for all f-Chrimson injected animals. The data is separated into the injected left (L) and non-injected right (R) cochlea and into animals with a positive oABR response (+) high threshold response (ht) and no oABR response (-) with up to 3 data points for the apex, mid and base turn per cochlea. For the density we processed non injected (n.i.) wildtype gerbil cochleae as a control group for both slicing types. All plots show the mean and 95% confidence interval and statistical comparison by Kruskal-Wallis test with Tukey-Kramer correction of multiple comparison (*** p<0.001, **p<0.01, *p<0.1).

oABRs were detectable in 34/48 (70 %) gerbils out of which 26 showed responses for stimuli below 10 µJ (54 %, average threshold 4.5 ± 2.6 µJ). These oABR values were in accordance to previously measured values in postnatally f-Chrimson injected gerbils (Huet *et al*, 2021; Thirumalai *et al*, 2025). Animals with a high oABR threshold (>10 µJ, ht) usually did not undergo further surgery and, if acquired, midbrain data was later excluded from the analysis, due to the transduction being too unreliable in these animals. oABR intensity thresholds of the included f-Chrimson animals were then compared to previous estimates in gerbils with other ChRs (Fig. 1B). They were higher than for oABR-positive, ChReef injected gerbils (0.28 ± 0.19 µJ, n=17, p<0.01, Kruskal wallis with multiple comparisons (Albrecht et al., companion study and (Alekseev *et al*, 2025)), and not significantly different from oABR-positive CatCh injected gerbils (4.59 ± 2.88 µJ, n=14, (Wrobel et al, 2018)).

At the end of the surgery the injected left (L) as well as the non-injected right (R) cochleae were extracted, fixed and processed for immunohistological analysis and quantification of cell survival and the transduction efficiency (see Fig. 1C). We performed either cross-modiolar (220 µm vibratome sections providing larger samples of SGNs and hair cells of individual turns) or mid-modiolar sections (25 µm cryosections near a central vertical plane assessing SGNs of all cochlear turns within one slice). We stained against parvalbumin (PV) for identifying SGNs and GFP to detect the eYFP tagged f-Chrimson construct (GFP^+^ SGNs). We then used a Cellpose model (Stringer *et al*, 2021) to segment the cells in the PV channel. We measured the GFP signal in SGNs and estimated the fraction of GFP^+^ SGNs (Fig. 1D) that marks a lower bound for the transduction efficiency as some transduced SGNs might have been lost over the time since AAV-injection. The injected ears of oABR positive, low-threshold animals showed 50 ± 24 % (mean ± std, n =39, ‘L+’) GFP^+^ SGNs. The right ear of these animals showed 21 ± 23 % GFP^+^ SGNs (n =37, ‘R+’). Animals with high oABR thresholds (>10 mW) had 32 ± 25 % GFP^+^ SGNs in the injected ear (n =14, ‘Lht’) and 20 ± 25 % GFP^+^ SGNs in the non-injected side (n =14, ‘Rht’). The high-threshold data split into some cochleae largely lacking f-Chrimson transduction while others had a fraction of GFP^+^ SGNs similar to the oABR-positive, low-threshold gerbils. oABR-negative animals had a low fraction of GFP^+^ SGNs in their injected ear (mean: 18 ± 15 %, n =33, ‘L-’) while their non-injected ears typically lacked GFP^+^ SGNs (0 ± 16%, n =38, ‘R-‘). The cell density was slightly reduced in the injected cochleae of oABR-positive animals (Fig. 1E, 20.9 ± 9.5 SGNs/10^4^ µm^2^, n =39) compared to their non-injected cochleae (28.4 ± 9.2 SGNs/10^4^ µm^2^, n=37), and left and right cochleae of non-injected gerbils (27.9 ±8.4 SGNs/10^4^ µm^2^, n =50, ‘n.i.’). For animals with a high oABR threshold, density for the injected (22.4 ± 11.2 SGNs/10^4^ µm^2^, n =13) and non-injected ears (29.7 ± 7.5 SGNs/10^4^ µm^2^, n =14) were not significantly different from the n.i. control animals. Also, no density differences were observed for oABR-negative animals in their injected (24.9 ± 9.3 SGNs/10^4^ µm^2^, n =33) and non-injected (28.6 ± 8.7 SGNs/10^4^ µm^2^, n =38) side.

### Recordings of multiunit activity in the inferior colliculus upon f-Chrimson-mediated SGN stimulation

In animals with low oABR thresholds (<10 µJ), we proceeded to recording from the auditory midbrain (Fig. 2A). Following a posterior craniotomy and stereotactically placed a 32-channel linear silicon MEA in the ICC for recording MU activity along its dorso-ventral tonotopic axis (low-high frequencies, surgical illustration in Fig. S2E-F). We verified our MEA placement by playing a set of 100 ms pure tones between 0.5 and 32 kHz and assigning each frequency a best electrode - the electrode with the lowest threshold for this frequency. We performed a linear regression of best electrode and tone frequency to extract the tonotopic slope of the ICC frequency map (Fig. 2B) and only animals with a slope in the range [3, 8] oct/mm were included in the analysis. The mean tonotopic slope of all included f-Chrimson animals was 5.5 ± 0.9 oct/mm (n = 25). Tonotopy is also illustrated by the mean best electrode estimates for low, mid and high frequency ranges (0-2, 2-8, 8-32 kHz, Fig. 2C). This dorso-ventral shift observed for acoustic stimulation of the apex, mid and base of cochlea, was also found for optogenetic stimulation with 50 or 200 µm optical fibers delivering of 1ms, 594 nm light pulses via apical, mid or basal cochleostomies (Fig. 2C, surgical illustration in Fig. S2B-D).

**Figure 2.**
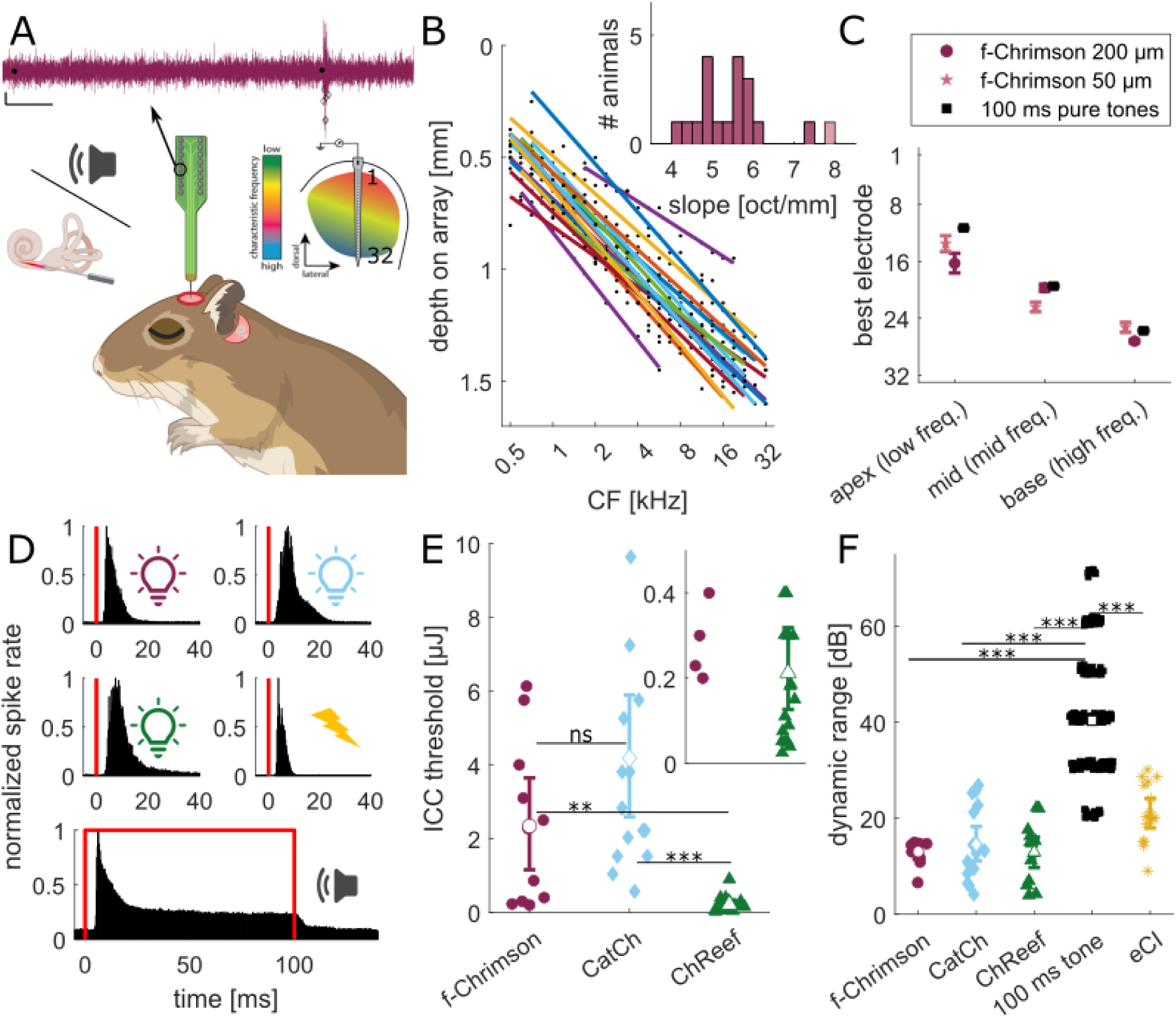
Inferior colliculus activity evoked by f-Chrimson-mediated optogenetic SGN stimulation in gerbils. **(A)** We placed a 32-channel electrode array in the tonotopically ordered ICC to record multiunit activity in response to acoustic and optical stimulation of the cochlea. Top: exemplary recordings from a single electrode in response to 0.6 µJ (1st) and 12.6 µJ (2nd) 1 ms, 594 nm stimulation (black dots are triggers, diamonds are detected spikes, scalebars are 100 ms and 10 µV) Bottom: Method illustration (Created in BioRender. Koert, E. (2026) https://BioRender.com/dmg8j8k). **(B)** The best electrode was determined at the beginning of each recording session as the electrode with the lowest response threshold for 100 ms pure tones (data points show position of the best electrode). A linear fit was used to determine the tonotopic slope. Each colored line represents one animal. The mean slope for all f-Chrimson animals was 5.5 ± 0.96 oct/mm. Animals with a slope ≤3 or ≥8 oct/mm were excluded from the analysis. **(C)** When placing the optical fiber in either the round window (base) or in mid- or apical cochleostomies the best electrode position shifts similar to the response to pure tones when separating the frequency range into apex (0.5-2 kHz), mid (2-8 kHz), and base (8-32 kHz). **(D)** PSTHs for 1 ms light pulses presented through the round window for f-Chrimson (32 µJ, 594 nm), CatCh (32 µJ, 488 nm, reanalyzed from (Dieter et al, 2019), and ChReef (1 µJ, 522 nm, reanalyzed from (Albrecht et al., companion study and (Alekseev et al, 2025)), a 0.1 ms biphasic pulse from the first electrode of a rodent eCI system in monopolar stimulation mode (100 µA, reanalyzed from (Dieter et al, 2019; Keppeler et al, 2020) and the responses to 100 ms pure tones between 1 and 16 kHz collected in the f-Chrimson injected gerbils. **(E)** ICC activation thresholds for the different opsins in injected gerbils measured with 200 µm fibers through the round window. The CatCh data is reanalyzed from (Dieter et al, 2019; Michael et al, 2023), the ChReef data is taken from (Albrecht et al., companion study and (Alekseev et al, 2025)). **(F)** Dynamic range as the intensity range between eliciting 10 and 90 % of the max. evoked spike rate at any electrode in dB rel. to the ICC threshold for each recording: significant differences were observed in comparison to acoustic stimulation, not between the bionic stimulation modalities. For the 100 ms pure tones only every 5^th^ value was plotted and small y-jitter ± 2 dB SPL was added to enhance visibility of the data points. All plots show the mean and 95% confidence interval and for statistics a Kruskal-Wallis test with Tukey-Kramer correction of multiple comparisons was performed (*** p<0.001, **p<0.01, *p<0.1).

For comparison to f-Chrimson-mediated responses, we also reanalysed ICC data from other studies, that included oCI animals that were stimulated with blue light pulses (1 ms, 488 nm, 200 µm fibers, adult cochlear AAV-CatCh-injection, (Dieter *et al*, 2019; Keppeler *et al*, 2020; Michael *et al*, 2023)) or with green light pulses (1 ms, 522 nm, 200 µm fibers, postnatal cochlear AAV-ChReef-injection, (Albrecht et al., companion study and (Alekseev *et al*, 2025)), as well as eCI animals that were stimulated with a rodent eCI system (Dieter *et al*, 2019; Keppeler *et al*, 2020). The temporal responses to each stimulus modality are different as demonstrated by the respective peristimulus time histograms (PSTHs, Fig. 2D; Fig. S4). For comparison between modalities for ‘single pulses’ we focused our analysis time window on the onset response of 2-15 ms for bionic stimulation and 5-18 ms for pure tone stimulation. Within these time windows we determined the ICC threshold as first intensity to reach a significant difference (d’>=1) when comparing the spike rate across 30 stimulus trials in the response time window to a baseline time window before trigger onset (-15 to -2 ms for optical and electrical stimulation and -18 to -5 ms for acoustic stimulation). The thresholds for f-Chrimson (mean ± std: 2.34 ± 2.20 µJ, n =11, oABR threshold <10mW) were similar to CatCh (4.18 ± 3.49 µJ, n =16, no selection for oABR threshold implied) and significantly higher compared to ChReef (0.21 ±0.22 µJ, n =17, oABR threshold <10mW). This threshold was independent of fibre diameter or position of cochleostomy (see Fig. S5). We estimated the output dynamic range as the difference in presented stimulus intensity relative to the activation threshold between reaching 10 and 90 % of the maximal evoked spike rate across all electrodes (see Fig. 2F). Here direct bionic SGN stimulation had a similar dynamic range across modalities (mean ± std: f-Chrimson 13.04 ± 2.51 dB (mW), n =11; CatCh 14.56 ± 7.52 dB (mW), n =16; ChReef 12.98 ± 6.18 dB (mW), n =13; eCI 21.27 ± 6.25 dB (µA), n =15) while the range for pure tones was significantly larger (40.39 ± 13.06 dB SPL, n=920 frequencies from 71 animals). An overview of the evoked spike rates reached for each stimulus modality is presented in Fig. S6. Not all recordings reached a saturation of the spike rate for the highest investigated intensity, especially for optogenetic stimulation.

Overall, these results showed that f-Chrimson mediated optogenetic SGN stimulation with orange light allows for activation of the auditory pathway up to at least the ICC. We observed a tonotopic organization of the responses in line with the cochlear position of optogenetic stimulation. The activation threshold was in the range of a few µJ (for animals with good oABR thresholds) and the dynamic range was comparable to other bionic means of SGN stimulation.

### f-Chrimson mediated optogenetic SGN stimulation elicits ICC activity with high temporal precision

In addition to the previously described “single pulses” (Fig. 2), we also presented 100 ms long “pulse trains” (32 µJ, 594 nm) at different stimulation rates. We compared these responses to data for 100 ms trains of optogenetic stimulation with CatCh (32 µJ, 488 nm, (Michael *et al*, 2023)) or ChReef (1 µJ [due to low activation threshold], 522 nm, (Albrecht et al., companion study and (Alekseev *et al*, 2025)) and to acoustic stimulation (0.1 ms clicks, 75 dB (SPL) (data from (Michael *et al*, 2023)). This data was used to investigate the ability of the neurons to follow higher stimulation rates which is a relevant parameter for encoding of real-live acoustic stimuli.

f-Chrimson-mediated SGN stimulation enabled ICC MUs to follow higher rates compared to CatCh and ChReef as exemplified for 100 Hz stimulation in the PSTHs (Fig. 3A) and raster plots for single electrodes for the different modalities (Fig. 3B-E). Responses to acoustic clicks also follow high stimulation rates.Even for very high rates, spiking was elicited throughout the full 100 ms of stimulation, while for optical stimulation only onset responses remained at these rates. When quantifying the spike rate of the responsive Mus as a function of stimulation rate across all the animals we found that the spike rate in the time window [25, 125] ms after stimulus onset was the highest for acoustic clicks (Fig. 3F). Spike rate continuously grew reaching a plateau around 300 Hz response spike rate from 400 Hz of stimulation onward. In contrast, f-Chrimson stimulation showed a peak spike rate of 230 Hz around 190 Hz stimulation rate. The spike rate decreased to around 60 Hz at higher rates of stimulation from 400 Hz onward. The other ChRs revealed a similar response shape but had much lower response rates with peak rates at lower stimulation rates compared to f-Chrimson (CatCh: 68 Hz peak rate for a 90 Hz of stimulation, ChReef: 65 Hz peak rate at 50 Hz of stimulation). Past the peak, the firing rate dropped to around 50 Hz for CatCh and around 25 Hz for ChReef before starting to slowly increase again for the higher rates.

**Figure 3.**
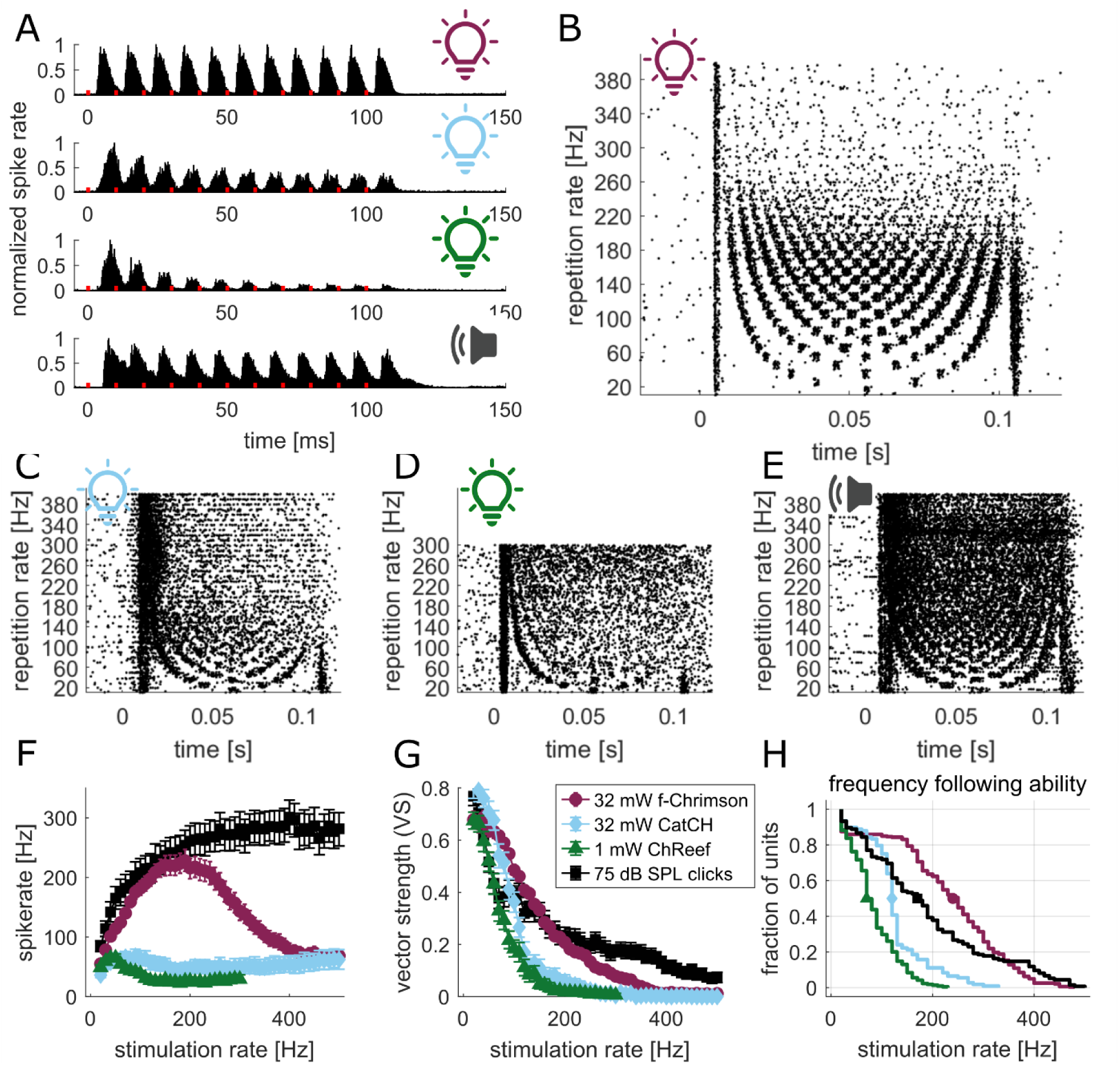
High temporal precision ICC activity evoked by f-Chrimson mediated SGN stimulation. We presented the gerbils with 100 ms stimulus trains of 1 ms light pulses (200 µm optical fiber via round window) or 0.1 ms acoustic clicks. Data for CatCh, ChReef and the clicks was reanalyzed from previously published recordings of (Alekseev et al, 2025; Michael et al, 2023, and Albrecht et al., companion study). **(A)** PSTH for the 100 Hz stimulation for 32 µJ, 594 nm light pulses for f-Chrimson (n=12), 32 µJ, 488 nm light pulses for CatCh (n=5), 1 µJ, 522 nm light pulses for ChReef (n=9) and 75 dB click stimuli in untreated gerbils (n=5). Stimulus timing is indicated by red marks on x-axis. **(B)** raster plot of spikes detected by an individual electrode in the center of activation for f-Chrimson recording over 30 presentations at each stimulation rate. **(C-E)** Raster plot as in B for CatCh, ChReef and clicks. **(F)** Average spike rate (across all responsive MUs) measured in the time window [25,125] ms after stimulus onset for each stimulation modalities as a function of stimulation rates. **(G)** Vector strength (VS) was calculated by aligning the detected spikes in the phase of the input stimulus train. mean and 95% confidence interval collected from 322 MUs in 12 animals for f-Chrimson, 108 MUs in 5 animals for CatCH, 220 MUs in 9 animals for ChReef and 156 MUs from 5 animals for clicks. **(H)** Fraction of responsive units that was able to follow up to a specific frequency with a significant VS (calculated with the Rayleigh test, alpha= 0.001). The markers show where 50 % of the units fail to follow the stimulation (240 Hz for f-Chrimson, 120 Hz for CatCH, 70 Hz for ChReef, 170 Hz for Clicks).

To evaluate temporal fidelity of ICC activity beyond spike rate, we calculated the vector strength (VS) of the detected spikes within the phase of the stimulation as a measure of temporal precision (Fig. 3G). We quantified how many MUs were able to respond with a significant VS for a given stimulation rate (Fig. 3H). VS of MUs that did not show significance in the Rayleigh test were set to 0 for calculating the average VS. First, we compared recordings of different stimulation modalities for strong stimuli (f-Chrimson: 32 µJ, 594 nm, CatCh: 32 µJ, 488 nm, ChReef: 1µJ, 522 nm, clicks: 75 dB SPL). With increasing stimulation rate, the number of units locking to the stimulation decreased for both optogenetic and acoustic stimulation. The f-Chrimson mediated responses showed a higher or equal mean VS compared to acoustic stimulation up to 200 Hz, for ChReef and CatCh the difference to acoustic was much smaller and a higher VS was only observed up to around 100 Hz. Beyond this the VS of CatCh and ChReef mediated stimulation declined to near zero (values below 0.025) around 200 Hz for ChReef and around 250 Hz for CatCh, while f-Chrimson stimulation retained non-zero VS until around 400 Hz. Responses to acoustic clicks plateaued at a slightly larger VS of around 0.06 for up to the maximal stimulation rate of 500 Hz. Fig. S7 shows the observed mean VS calculated not over all active MUs as in Fig. 3G but only over those that were still able to follow each frequency to illustrate the temporal precision in the remaining signal. When estimating the number of responsive MUs that were able to faithfully respond to increasing stimulation rates, f-Chrimson outperforms the other modalities. Half of the units still responded with a significant VS (Rayleigh test, see methods) at 240 Hz, while for ChReef, CatCh, and Click stimuli this occurred at 70, 120, and 170 Hz respectively. Around 350 Hz of stimulation we reach the point where the number of units that can follow the input of f-Chrimson falls below the acoustic stimulation.

In addition, we obtained a data set with lower laser intensities (16 µJ and 8 µJ) for f-Chrimson and CatCh (Fig. EV 1). f-Chrimson-mediated responses to 16 µJ showed similar or even slightly larger spike rates and VS compared to the stimulation at 32 µJ. We observed a reduction in spike rate at 8 µJ yet it still peaked at the same stimulation rate of around 200 Hz (32 µJ: 228 Hz response spike rate for 190 Hz stimulation; 16 µJ: 258 Hz for 250 Hz stimulation; 8 µJ: 96 Hz for 200 Hz stimulation). VS was smaller for 8 µJ than for 32 µJ at low stimulation rates and then reached the same or even higher VS for stimulation rates above 100 Hz. The number of MUs that were able to follow the stimulation rate was largest for 16 µJ (50 % were 240, 300 and 200 Hz for 32, 16 and 8 µJ).

**Figure EV 1.**
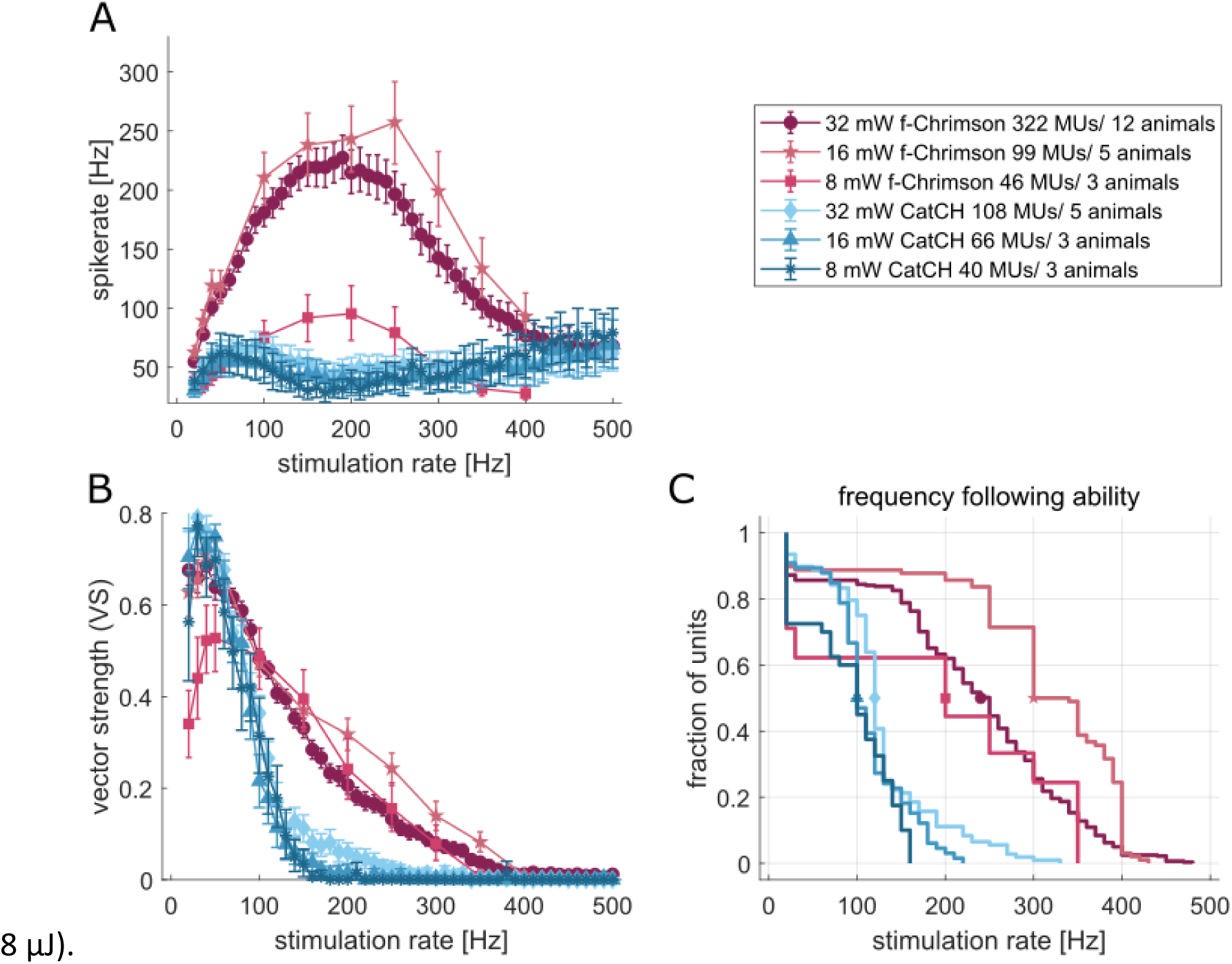
Reducing stimulus intensity decreases the spike rate but not the temporal fidelity of ICC activity. **(A)** Spike rate in response to different stimulation rates measured in responsive electrodes (MUs) in the time window [25,125] ms after stimulus onset for 1 ms pulses at 8,16 and 32 mW in f-Chrimson and CatCh (Michael *et al*, 2023) animals. **(B)** The vector strength (VS) was calculated by aligning the detected spikes in the phase of the input stimulus train. The Rayleigh criterion was used to test for significance and if no significance was reached, the VS was set to 0 for that rate in this MU. **(C)** Fraction of investigated MUs that was able to follow up to a specific frequency with a significant VS. The markers stress the point where 50 % of the units fail to follow the input stimulation rate (f-Chrimson: 240, 300 and 200 Hz; CatCh: 120, 100,100 Hz for 32,16 and 8 mW, respectively). The plots in A and B show mean and 95% confidence interval.

Finally, we separated the MUs responsive to f-Chrimson-mediated SGN stimulation depending on their position within the ICC. For this we assigned each electrode a characteristic frequency based on their responses during the initial acoustic recordings. We then separated the data at 2 and 8 kHz into low, mid and high frequency ranges of ICC neurons that receive input driven by optogenetic stimulation of the apical, mid- or basal cochlea, respectively. The optical stimulation was achieved using a 200 µm fiber placed at the round window such that it projected light along the modiolar axis and broadly activated SGNs across frequency (tonotopic) ranges. We illustrate the stimulation configuration by an X-ray tomography of a sham fiber insertion (Fig. EV 2A). We consider light to reach highest irradiance in SGNs of the basal high frequency cochlea and lower irradiance for mid-cochlear and apical SGNs. We compared the results for f-Chrimson-mediated SGN stimulation (32 and 16 µJ) to 75 dB acoustic clicks (∼30 dB above threshold). As expected, recordings with optical stimulation showed higher spike rates for high frequency MUs compared to the mid- and low-frequency ranges for both stimulus intensities. While no very clear differences in VS were found between the MUs of the frequency ranges in recordings with 32 µJ, high frequency MUs showed larger VS than mid- and low-frequency MUs in the 16 µJ recordings. The stimulation rates where 50% of the units failed to follow the stimulation were 210, 210 and 270 Hz for low, mid and high frequencies in the 32 µJ recordings and 250, 250 and 380 Hz in the 16 µJ recordings. As expected for the spectral content of the acoustic clicks, MUs of the mid frequency regions had the highest spike rate, while VS was similar between all groups. The 50 % following rates were 130, 180 and 190 Hz for MUs of the low, mid- and high frequency ranges.

**Figure EV 2.**
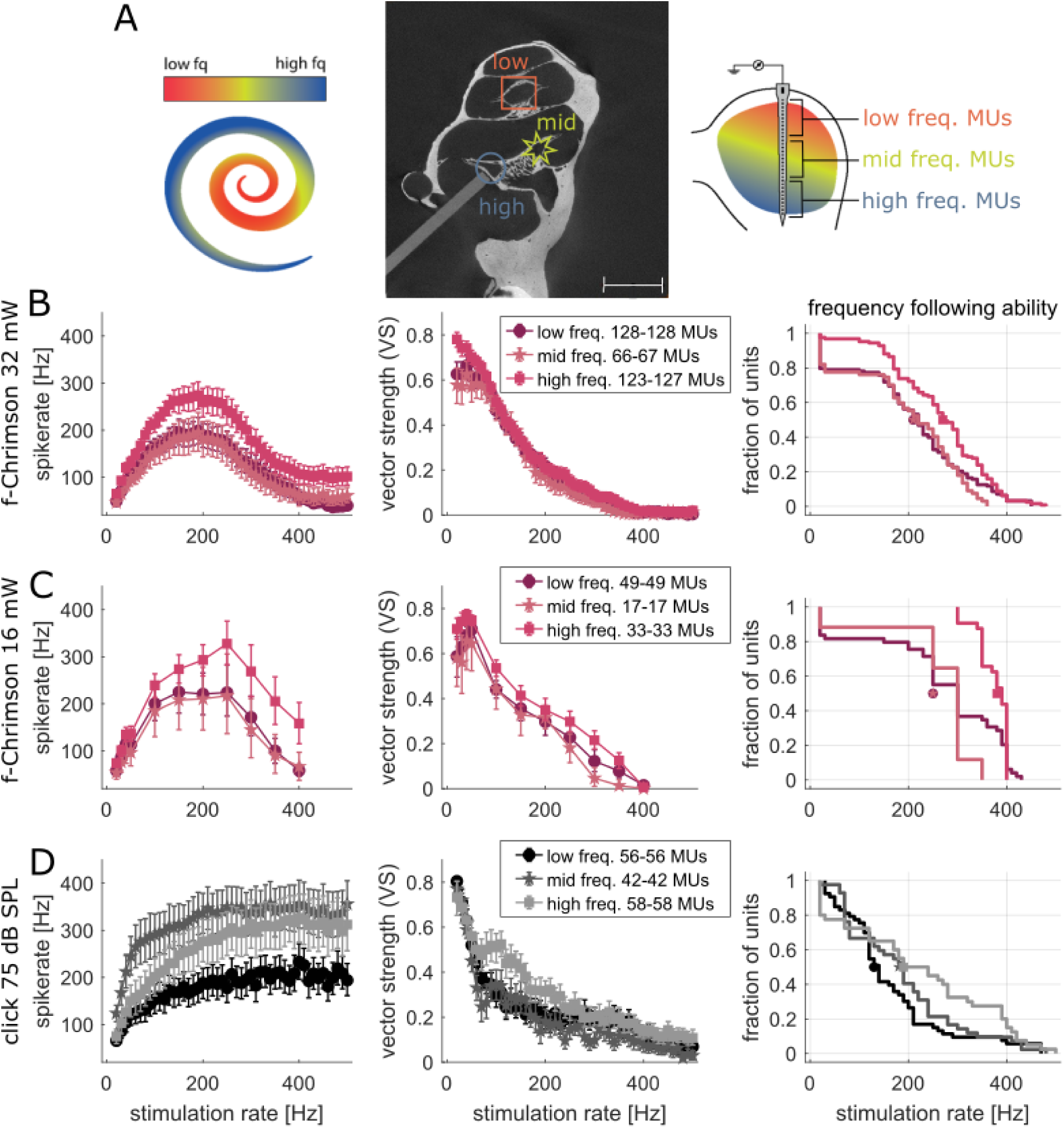
Separating the MUs into three tonotopic ranges shows that the temporal fidelity of f-Chrimson-mediated optogenetic stimulation depends on proximity to the stimulating fiber. We separated the f-Chrimson data into 3 groups depending on what best frequency was assigned to the responsive electrode in preceding recordings with pure tone stimulation. **(A)** Visualization of the method using an X-ray tomography of a gerbil cochlea with a 200 µm fiber implanted into round window (RW)). The fiber end is directly next to the base turn of the Rosenthal’s canal, which is where the high frequency SGNs are located that drive the high frequency area of the ICC. **(B)** Spike rate, vector strength (VS) and ability of neurons to follow up to specific stimulation rates for the f-Chrimson MUs stimulated with 32 µJ. MUs in the high frequency area of the ICC, getting input from the basal cochlea regions have a higher spike rate and frequency following ability. The number of responsive MU varies for different stimulation rates (range given in the legend) since different stimulation protocols were used for different animals. **(C)** Same plots as in B for 16 µJ stimulation. **(D)** The same separation of MUs was performed for the 75 dB click stimulation reanalyzed from (Michael et al, 2023). The VS and frequency following ability seems similar in all three ICC areas, while the spike rate is highest in the mid frequency range.

In conclusion, f-Chrimson mediated optogenetic SGN stimulation enables temporally precise ICC responses for high stimulation rates outperforming previously published opsins. Lowering the light intensity does not negatively affect the temporal precision as long as enough spikes are generated. The ability to respond well to high stimulation rates seems to depend on the irradiance achieved at the target SGN.

### The spatial precision of f-Chrimson modulated stimulation outperforms electrical stimulation

Previous recordings of ICC MU activity indicated that CatCh-mediated blue light optogenetic SGN stimulation achieves higher spectral selectivity than electrical stimulation (Dieter *et al*, 2019, 2020; Keppeler *et al*, 2020). Here, we investigated the spectral selectivity of f-Chrimson mediated optogenetic SGN stimulation using orange light delivered by 200 µm (as in (Dieter *et al*, 2019) and also by thinner (50 µm) optical fibers and evaluated how wavelength and emitter size might affect the cochlear spread of excitation (SoE). Fibers were placed into the round window (base) or into cochleostomies at the apex or mid-turn of the cochlea and ICC MU activity was recorded in response to 1 ms pulses varying in radiant energy.

We then measured the SoE for these recordings as the distance between the highest and lowest electrode in the ICC that was above threshold (d’ = 1 when comparing the response spike rate with the baseline before trigger) at different levels of evoked MU activity (Fig. 4A). For a robust comparison, we re-analyzed previously published data for CatCh-mediated optogenetic and electrical SGN stimulation from the raw data of (Dieter et al, 2019; Keppeler et al, 2020) using identical scripts and analysis time windows across modalities. Responses to pure tones were pooled across tonotopy recordings from all animals of the past and present study. A visual comparison of the SoE is provided in Fig. S8 that shows evoked spike rate heatmaps within the same intensity range for all stimulation modalities. We note that the present analysis differs from the previous one (Dieter et al, 2019; Michael et al, 2023) that used different time windows adjusted to the PSTH of each modality and measured SoE at different d’ levels while only looking at the best electrode. For our approach, we decided to measure the SoE whenever any electrode reaches a new evoked spike rate level instead. This was motivated by the fact that the area of highest MU activity tends to shift along the MEA especially for electric and optical stimulation (See Fig. S9). For consistency with the old publications we ran also ran the SoE analysis once using the fully d’ based analysis with different time-windows for each modality and keeping to the best electrode while reading out the intensity levels (see Fig. S10).

**Figure 4.**
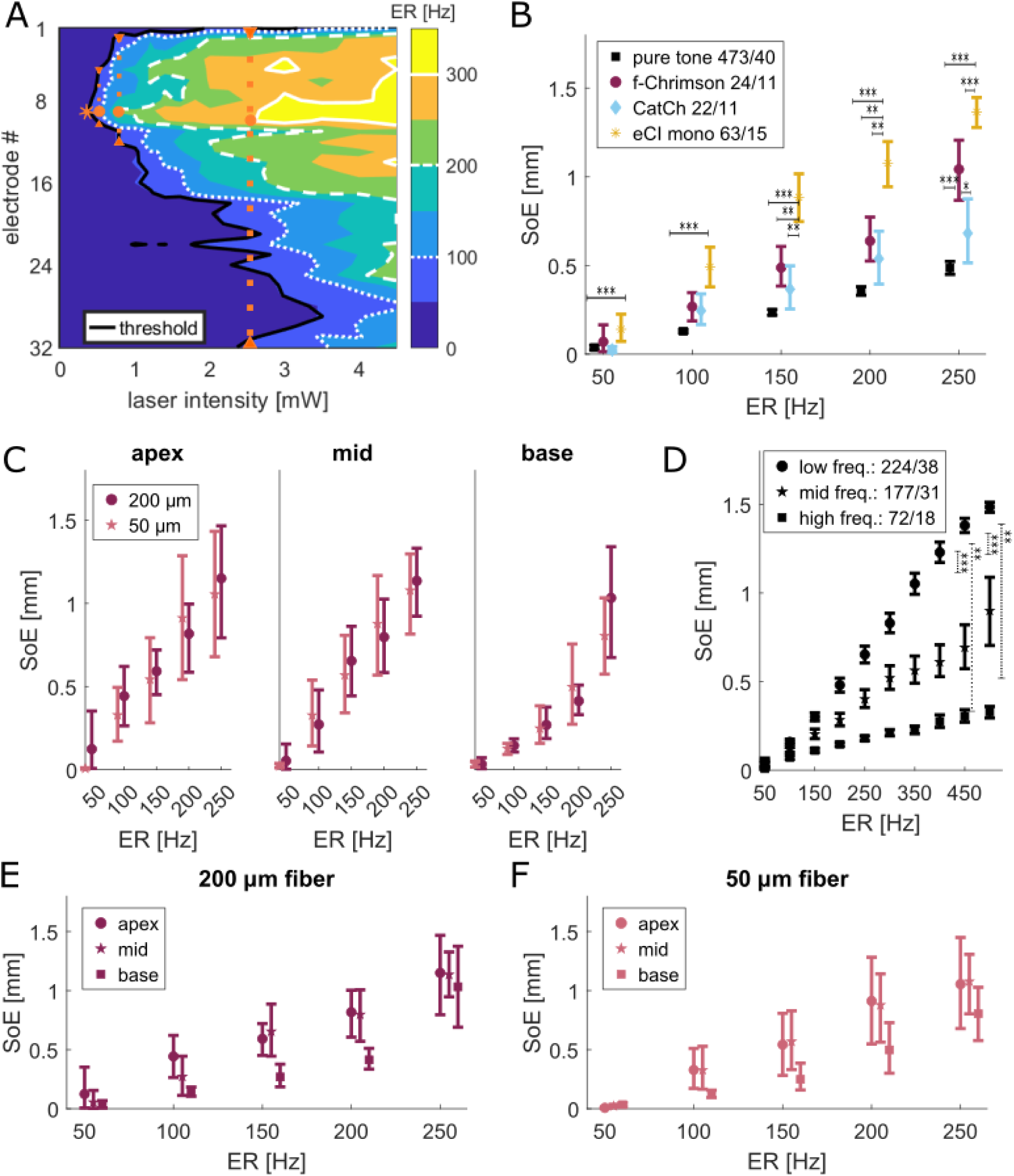
Spectral selectivity of f-Chrimson mediated optogenetic SGN stimulation. We compared ICC MU responses to 1 ms, 594 nm light pulses to those elicited by 100 ms pure tones and electrical pulses. To evaluate cochlear spread of excitation (SoE), we measured the number of electrodes that reached threshold (baseline d’ > 1) at the intensity when a specific evoked spike rate (ER) was reached using only the onset time window (2-15 ms for bionic stimulation, 5-18 for acoustic stimulation). **(A)** Spatial tuning curve for an exemplary recording of MU activity elicited by light delivered from a 200 µm optical fiber via an apical cochleostomy illustrating the SoE measurement as the width (number of active electrodes) of the threshold iso-contourline (d’=1, black line) at the first occurrence of the evoked ER levels of 100, 200 and 300 Hz. In this example we have 4.5, 10.4 and 31.0 active electrodes for the different levels. To get from the number of active electrodes to the spread in mm, we multiplied it with the electrode pitch of 50 µm. **(B)** Mean SoE for f-Chrimson (all 200 µm fibers in apex, mid, or base) and reanalyzed CatCh (all 200 µm fibers in apex, mid, or base) and eCI (all emitters) data (Dieter et al, 2019; Keppeler et al, 2020), and to pure tone responses from all animals (old datasets and new recordings, full frequency range). The numbers in the legend show number of MUs/animals. Only recordings that reached a ER of 250 Hz were included. If multiple recordings existed for a diameter and cochlea position the one with the lowest spread at 100 Hz was chosen. **(C)** Splitting f-Chrimson data by fiber diameter and stimulation position within the cochlea: the number of recordings was 8, 9, 8 for the 200 µm fibers and 5, 8, 8 for the 50 µm fibers for apex, mid and base respectively. No significant differences were found when comparing SoE between fiber diameters. **(D)** Separating SoEs for pure tone recordings into ICC frequency areas (low frequencies < 2 kHz corresponding to apical stimulation, mid frequencies: 2-8 kHz, high frequencies > 8 kHz corresponding to basal stimulation) over a larger ER range. Low frequencies have a larger SoE than mid and high frequencies especially for high ERs. **(E-F)** SoE does notdiffer significantly across stimulation sites for f-Chrimson-mediated SGN stimulation for 200 µm **(E)** or 50 µm **(F)** fibers. All plots show the mean and 95% CI. We fitted a repeated measure model with Tukey-Kramer correction of multiple comparison (*** p<0.001, **p<0.01, *p<0.1) to test the differences between groups.

When pooling all recordings across cochlea turns (Fig. 4.B), f-Chrimson- and CatCh-mediated optogenetic stimulation had quite similar mean SoEs with slightly larger SoE for f-Chrimson at the highest investigated spike rate (250 Hz). Both optogenetic methods activate slightly larger areas then the pure tone stimulation, but again only around 250 Hz the SoE differs significantly across the modalities. In contrast, the SoE for electrical stimulation was significantly larger than pure tone stimuli already around threshold (50-100 Hz evoked spike rate). For higher levels of MU activity, it was also significantly larger than the optogenetic stimulation. Since the maximum possible ICC spread is 1.55 mm (pitch between electrode 1 and 32) the measured spread saturated around 1.5 mm. For 250 Hz most eCI recordings reached this full ICC activation and some of the f-Chrimson recordings got close as well, causing the difference between the two groups to be less significant. CatCh and pure tone recordings remained far from activating the full ICC even at 250 Hz.

As a next analysis step, we separated SoEs obtained with f-Chrimson by fiber diameter and cochlear position (Fig. 4C, E and F). We found no significant differences between the diameters within a cochlear position or for one fibre diameter between cochlear positions. There was a trend toward smaller SoE for stimulation of the cochlear base. We also grouped acoustic ICC MU responses into frequency ranges by splitting the dataset at 2 and 8 kHz (Fig. 4D). Stimulation of the cochlea with low frequency tones tended to cause larger SoE compared to mid and high frequency tones for similar levels of activation.

To supplement our *in-vivo* findings we performed *in-silico* modelling of fibers at different cochlea positions following our previously described framework (Dieter *et al*, 2019; Khurana *et al*, 2022)(Fig. S11). The geometry for the model was taken from X-ray tomographies of cochleae with fibers glued in position during training surgeries on sacrificed animals. Some of these fibres were positioned well, pointing towards Rosenthal’s canal (RC), while others were off-target allowing for a comparison of the effects of fiber placement. The simulated irradiance was measured and the SoE determined as described in the methods. The default simulation used a diameter of 200 µm, a wavelength of 594 nm and a numerical aperture of 0.5. The main factor affecting the spread of light in the model was the fibre placement. Changing the diameter of the fibres or wavelength had no effects and changing the numerical aperture showed small changes, with narrower light beams slightly reducing the spread of light for fibres that were pointed towards the RC, but the differences were not statistically significant in this dataset.

In summary, our ICC MU recordings of f-Chrimson-mediated SGN stimulation corroborates the previously demonstrated improved spectral selectivity compared with electric stimulation for longer wavelength light. We did not observe greater spectral selectivity upon using narrower fibers.

### Short and weak light pulses enable energy-efficient and spatially confined neural responses

After this in-depth analysis of spectral and temporal ICC response properties elicited by 1 ms pulses or trains of 1 ms pulses we tested different pulse durations at varying intensities to optimize stimulus conditions for f-Chrimson-mediated optogenetic SGN stimulation (Fig. 5). Longer pulse durations and higher light intensities drove SGNs more strongly causing higher ICC MU activity (Fig. 5B) as previously observed with CatCh and Chronos (Mittring *et al*, 2023; Michael *et al*, 2023). When comparing spike rate distributions after stimulus onset with the baseline before the trigger there was a general trend for higher d’ values with long, high intensity stimuli. The mean highest d’ values (max d’) reached for the best electrode in 16 animals are plotted in Fig. 5C, and the individual max d’ values are presented in (Fig. S12A, B). Stimulus combination that did not reach a d’ of 1 were set to 0. When dividing the mean max d’ values by the applied radiant energy we found an advantage of shorter and weaker light pulses for efficient coding (Fig. 5D). For higher radiant flux (32 mW) we could use shorter pulses down to 0.1 ms to activate the ICC, while lower intensity pulses require longer durations. The highest max d’/radiant energy ratio was achieved with a 0.7 ms pulse at 2 mW. Yet, 0.5 to 1 ms at 2 mW as well as 0.4 ms at 4 mW stimuli also resulted in efficient stimulation. When counting the number of active electrodes that reach a d’ value above 1 for each stimulus condition as a proxy of SoE, the number of active electrodes increased with pulse duration and intensity (see Fig. S12C and D). Here, the sweet spot of reaching a high response strength at the best electrode with minimal SOE seems to be in the same range of 0.6-1 ms pulses close to the ICC activation threshold at 2 mW.

**Figure 5.**
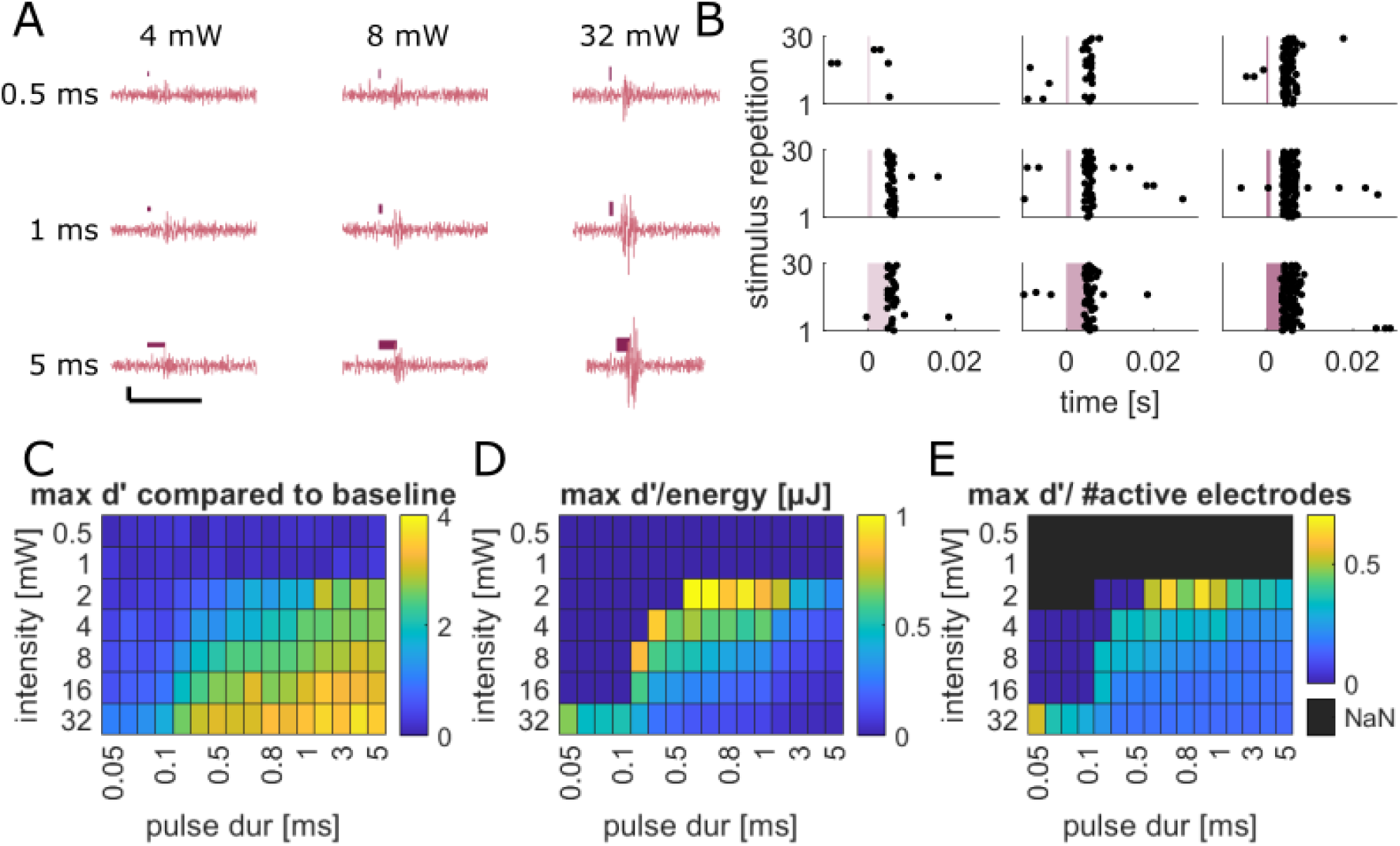
Short pulses at low light intensities cause efficient and spectrally selective SGN stimulation. **(A)** Exemplary filtered raw data traces from one ICC electrode in response to pulses of different duration (0.5, 1, 5 ms) and intensities (4, 8, 32 mW @ 495 nm) measured in response to f-Chrimson-mediated SGN stimulation with a 200 µm fiber in the round window. The dark red bars indicate the stimulus timing and intensity. The scalebar shows 20 ms and 20 µV. **(B)** Raster plots for the same recording as in A showing the detected spikes over all 30 stimulus repetitions. **(C)** Maximum d’ (max d’) value reached at any electrode when comparing the spike rate in the full f-Chrimson specific response time window [2,18] ms with the baseline before the trigger [-18,-2] ms (mean results from 16 recordings from 16 animals including all fiber positions and diameters). **(D)** Ratio of max d’ and pulse radiant energy. Stimulus combinations that did not reach a d’ value of 1 were set to 0 (lack of activation). **(E)** We divided the max d’ values from C by the mean number of active electrodes for each stimulus condition independent of their position within the IC.

## Discussion

In this study we scrutinized the utility of f-Chrimson for bionic encoding of frequency, time and intensity information by recording MU ICC-activity evoked by SGN stimulation. We demonstrate modest light requirements (threshold below 5 µJ) similar to those reported for CatCh (Dieter *et al*, 2019) but above those of ChReef (Alekseev *et al*, 2025). Owing to its fast kinetics, f-Chrimson offered excellent temporal response dynamics to light pulses with sharp on- and off-responses of neural activity. The ability of ICC activity to follow f-Chrimson optogenetic SGN-stimulation was superior to that found with the slower ChRs CatCh and ChReef and even exceeded that observed with acoustic clicks. The spectral spread of cochlear excitation (SoE) was comparable to the one previously measured with CatCh, lower than for monopolar electrical stimulation, but greater than for pure tone stimulation. We did not find SoE differences for 50 and 200 µm fibers placed at apical, mid and basal cochleostomies. The dynamic range of ICC-activity to single pulses compared well to the other opsins and electrical stimulation but was substantially lower than for acoustic stimulation as expected for direct neural stimulation. Postnatal gene therapy was reliable and provides an upper bound for the efficiency of optogenetic stimulation yet also illustrated the risk of neural loss in cases with high virus titers. In summary, this study supports the view that improved spectral coding by optogenetic stimulation can go along with good temporal coding provided fast ChR kinetics.

### Strengths and challenges of comparing optogenetic SGN stimulation across studies

Here, we preclinically evaluated the utility of f-Chrimson, a candidate ChR tailored to optogenetic hearing restoration (Mager *et al*, 2018; Bali *et al*, 2021; Huet *et al*, 2021; Bali *et al*, 2022; Thirumalai *et al*, 2025) at the level of the ICC which is a central auditory hub representing temporal, spectral and intensity information (Drotos & Roberts, 2024). For a comparative approach, we included ICC raw data previously recorded in the same condition in response to optogenetic, acoustic and electrical SGN stimulation (Dieter *et al*, 2019; Keppeler *et al*, 2020; Michael *et al*, 2023; Alekseev *et al*, 2025). This allowed us to apply the same analysis functions to all data to reduce methodological biases in the comparison of data for different stimulation modalities while also introducing more automated steps that further increase reproducibility and reduce the amount of human input necessary compared to earlier work (Dieter *et al*, 2019). By applying our new scripts to already acquired data we also circumvented the need for additional animal experiments and still gained new insights and a larger number of recordings for statistical comparison. The study also addresses reproducibility of auditory systems physiology as the recordings were performed by different experimenters over a decade. Despite all efforts to maintain laboratory standards e.g. on laser fiber and recording electrode placement, individual surgical experience might impact results and hardware at the setups as well as the stimulation and recording software underwent changes over time. Finally, stimuli were not entirely identical across studies, leading to different numbers of recordings available for the different stimulus modalities. These and other caveats present limitations of this study (see also below).

### Assessing the temporal fidelity of optogenetic coding of auditory information with a fast ChR

Optogenetic SGN stimulation bears great potential for improving spectral coding (Dieter *et al*, 2019, 2020; Keppeler *et al*, 2020; Alekseev *et al*, 2025), this study). However, given the millisecond-closing kinetics of most ChRs, the question remained whether this is traded in for poorer temporal coding. Good temporal coding underlies auditory functions such as sound source localization and is thought to be critical for speech understanding. Electrical CIs typically encode temporal fluctuations of the sound envelope rather than temporal fine structure. It seems that a stimulation rate of 500 Hz suffices speech understanding and even much lower rates (<100 Hz) have been shown to support speech understanding (Friesen *et al*, 2005).

Significant VS of optogenetically evoked firing of single SGNs was found to be limited to stimulation rates of 150-300 Hz even for fast ChRs such as Chronos (Keppeler *et al*, 2018) and f-Chrimson (Bali *et al*, 2021). The present study employed the central and more comprehensive perspective of the inferior colliculus in the auditory midbrain for evaluating the temporal fidelity of f-Chrimson-mediated optogenetic SGN stimulation. The data indicate that the ICC responds to high rates (up to 400 Hz) of f-Chrimson-mediated SGN stimulation with a temporal precision at least as high as found for trains of acoustic clicks. For stimulation rates up to 200 Hz, ICC MU spiking rates for SGN stimulation by 100 ms trains of 1 ms, 32 mW, 594 nm pulses closely resembled those for 100 ms, 75 dB SPL click trains. For higher stimulation rates, the ICC spiking rate dropped for f-Chrimson-mediated SGN stimulation while the spike rate plateaued for acoustic clicks. Yet, there were still responses to f-Chrimson-mediated SGN stimulation up to nearly 500 Hz. Potential causes of the drop in ICC spiking rate for higher rates of f-Chrimson-mediated SGN stimulation include a i) depolarization block of SGNs and ii) sideband inhibition at or beyond the level of the cochlear nucleus. Comparison to previously tested slower ChRs like CatCh and ChReef indicates superior temporal coding with f-Chrimson. A spike rate reduction was also evident for CatCh and ChReef albeit at much lower stimulation frequencies and to lower mean values.

f-Chrimson-mediated stimulation of SGNs yielded the highest number of MUs during high-frequency stimulation and showed enhanced VS compared to other ChRs and acoustic click stimulation (up to ∼230 Hz). Such high temporal precision is desirable for bionic sound encoding. eCI stimulation of SGNs can evoke even more time-locked responses at low pulse rates due to direct neuronal activation (Zeng *et al*, 2008; Wilson & Dorman, 2008). However, strictly synchronous activation is not necessarily optimal for information encoding, and the deliberate introduction of temporal jitter has been proposed to enhance neural coding by reducing excessive synchrony (Bruce *et al*, 1999; Carney, 2018). Despite high temporal precision at low rates, eCI stimulation remains limited in encoding temporal fine structure, particularly at higher frequencies (Shannon *et al*, 1995; Friesen *et al*, 2005). In this context, the increased VS achieved with f-Chrimson—approaching acoustic stimulation—represents a substantial improvement over previous optogenetic approaches and may support more flexible encoding strategies by balancing temporal precision with physiologically relevant variability.

The present study also tested whether temporal fidelity of f-Chrimson-mediated stimulation of SGNs can also be achieved at lower stimulus intensities. Halving radiant energy per pulse (16 µJ) did not reduce spike rate and even increased the number of MUs following high frequencies with a similar overall VS. While strongly decreasing the spike rate, further reduction to 8 µJ still allowed MUs to follow up to and above 400 Hz without shifting the peak spike rate around 200 Hz of stimulation. Interestingly, the fraction of MUs following high frequencies for 8µJ was only slightly smaller than for the 32 µJ. Yet, VS was smaller especially for low stimulation rates which might be attributed to the spike rate dependency of VS calculation and statistical evaluation (Rayleigh statistics takes both VS and spike count into account and set MU activity to a VS of 0 if the Rayleigh criterion was not fulfilled). We tried to circumvent this analysis bias in Fig. S7, where we plotted only the mean over units that fulfilled the Rayleigh criterion. While the 8 µJ stimulation still has a weaker VS at 10 HZ, it performs as well or even better than the higher stimulation intensities, when we only look at the units able to follow the stimulation. So, while with weaker light pulses, the spike probability goes down and more units fail to respond with high enough spike rates to reach significant VS levels, those that still respond to the stimulus do carry very precise temporal information.

Finally, temporal fidelity was greater in ICC areas close to the stimulating fiber than in those further away. Therefore, we hypothesize that temporal coding might improve when using more translational multichannel oCI inserted into the scala tympany. There, emitters are placed in close proximity to the spiral ganglion in RC along the tonotopic axis (Keppeler *et al*, 2020)and companion paper).

### Lessons learnt on spectral selectivity of optogenetic SGN stimulation

The previous observation of near physiological spectral selectivity for optogenetic stimulation (Dieter *et al*, 2019) held for our f-Chrimson data: no significant difference in comparison to the data obtained with CatCh and lower spread of excitation compared to electric stimulation. Here, we also probed the impact of emitter size using 50 and 200 µm optical fibers. Moreover, we performed simulations of light propagation for different stimulation parameters to gain further insight into what influences the spectral selectivity.

We expected to find a smaller SoE for the 50 µm fibers, which, however, was not consistently observed experimentally. Compared to the 200 µm fibers, it was more difficult to place and orient the 50 µm fibers at the cochleostomies to reliably evoke ICC activity. Moreover, computer simulations based on X-ray tomographies of sham fiber insertions at typical positions did not predict a systematic difference in the relative SoE for the same fiber position for fiber diameters of 200 and 50 µm. Instead, 3D modeling cautioned that emitter placement is critical for rays to hit the target: SGN somata in the RC. This targeting can readily be off with little or no exposure of the SGNs. Fiber positioning seemed unreliable such that the smaller light beam of the 50 µm fiber might have more often missed the target rather than to improve spectral selectivity. In the future it will be of interest to repeat the comparison in a more reproducible manner with scala tympani-inserted oCIs of different emitter diameters and shaped to facilitate the correct emitter orientation. Indeed, previous simulations of oCIs in the human cochlea revealed smaller SoE of waveguide emitters over LEDs and smaller over larger numerical apertures (Khurana *et al*, 2022). However, the above-mentioned challenge of missing the target for narrow beams, e.g. in case of oCI rotation was clearly indicated in this study that argued for balancing spectral selectivity and robust SGN stimulation when choosing the emitter.

So far computational studies have focused on either blue or red light applied to the cochlea. Here, we chose to compare blue- and red-light propagation in the gerbil cochlea, expecting differences related to less scattering of longer wavelength light as shown in previous work (e.g. Yizhar *et al*, 2011). However, we did not significant differences among these wavelengths within the limits of our modelling effort. This is in line with the fact that our comparison of ICC analysis of cochlear SoE for blue-light activated CatCh and the red-light activated f-Chrimson that did not reveal significant differences but was prone to experimental variance (see above). Nonetheless, our *in-silico* and *in-vivo* findings indicate that the choice of wavelength is less of a concern for spectral selectivity but wavelength differences need to be kept in mind in future studies on phototoxicity, which remains a highly relevant topic to investigate in the field of optogenetic stimulation of the cochlea.

When analyzing ICC data across modalities, we noticed that the peak of activation shifts along the tonotopic axis as a function of stimulus intensity. This was most pronounced for electric stimulation and less obvious for acoustic or optogenetic stimulation of the cochlea (see Fig. S11). We addressed that by allowing our SOE analysis to register a different best electrode for each intensity level to avoid overestimating the SoE for the eCI. However, this more conservative approach did not change the significance of the improved spectral selectivity of oCI compared to eCI stimulation. The shift of ICC activity evoked by eCI might reflect the current spread within the cochlea and explain shifts in perceived pitch with increasing loudness observed by CI patients (Umat *et al*, 2006). Our finding that this phenomenon is less for the optogenetic stimulation for similar levels of ICC activity informs the design of oCI sound coding strategies (Khurana *et al*, 2023) for which frequency can be mapped to one emitter for a larger range of sound intensities. Of course, this would need to be tested with a behavior setup for pitch detection in rodents or in a psychoacoustic test with oCI users in the future.

### Evaluating the potential of f-Chrimson as candidate for future optogenetic hearing restoration

Among currently available ChRs, f-Chrimson remains one of the strongest candidates for clinical optogenetic hearing restoration, especially when judged by the combined requirements of red-shifted activation, fast kinetics, usable light sensitivity, and an emerging preclinical safety package. Mager et al. (Mager *et al*, 2018) established f-Chrimson as a Chrimson variant with accelerated closing kinetics while preserving longer-wavelength activation, robust membrane expression, efficient AAV-mediated expression in spiral ganglion neurons (SGNs), low optical auditory brainstem response thresholds, and auditory-pathway responses following stimulation up to at least 200 Hz. This temporal fidelity is important because an oCI will likely rely on coding strategies that trade the extreme synchrony of electrical stimulation for more spatially confined, yet temporally precise optical activation. Follow-up work supported this view by showing that red-light ultrafast Chrimson variants can drive SGNs with near-physiological temporal fidelity; f-Chrimson offered better light sensitivity than vf-Chrimson, whereas vf-Chrimson improved kinetics, highlighting f-Chrimson as a pragmatic balance between optical power demand and timing (Bali *et al*, 2021). Its red-shifted activation is another translational advantage because 594 nm light is expected to be less phototoxic than blue light and better suited for cochlear illumination. Yet, a still deeper red action spectrum would be preferable for optimal compatibility with efficient red laser diodes and waveguide systems, which perform best at longer wavelengths. The pulse-duration data are also encouraging: activation with sub-millisecond pulses supports coding strategies using brief, energy-efficient stimulation epochs rather than long pulses, which should help constrain thermal load and total optical dose.

The main limitation is that f-Chrimson is no longer the most light-sensitive benchmark; newer candidates such as ChReef show substantially lower oABR energy thresholds, meaning that f-Chrimson may impose tighter demands on emitter output, which is challenging especially for high-rate multichannel stimulation. On the safety side, however, data on f-Chrimson is comparatively advanced: long-term, up to 24-month, AAV-mediated f-Chrimson expression in mouse SGNs after a single cochlear dose has been reported (Bali *et al*, 2022), supporting durability and gene-safety feasibility. Clinical derisking was further strengthened by removing the fluorescent protein tag and replaced it with a human Kir2.1 trafficking sequence; although tag removal reduced photocurrents and required optimization, f-Chrimson-TSKir2.1 restored optogenetically evoked auditory activity with only mild neural loss, making the construct more suitable for translation (Zerche *et al*, 2023). The remaining development path should therefore focus less on proving that f-Chrimson works and more on optimizing the full therapeutic system: matching waveguide/emitter wavelength to opsin sensitivity, reducing optical thresholds, and performing careful AAV titer-optimization studies with oABRs and histology to minimize SGN loss and off-target hair-cell activation.

### Limitations of the present study and future objectives

A number of limitations should be considered when interpreting the present results, particularly with respect to how spread of excitation (SoE) and temporal coding were assessed. First, our analysis relies on multi-unit (MU) activity in the ICC, rather than single-unit (SU) recordings. While MU recordings enable robust sampling across larger neuronal populations, they inherently blur the contribution of individual neurons and introduce additional noise, limiting precise conclusions about neural coding fidelity and potentially obscuring fine differences between stimulation modalities. This constraint is especially relevant when comparing optogenetic and electrical stimulation, where subtle differences in synchrony and recruitment patterns may be critical.

Second, the estimation of SoE itself is influenced by the anatomical and functional organization of the ICC. Our approach assumes a relatively uniform mapping between recording sites and cochlear frequency regions, yet isofrequency bands vary in width and representation across the ICC. As illustrated by the spread of pure tone responses (Fig. 4D), different cochlear regions are not equally represented, which may bias apparent resolution. Similarly, comparisons to eCIs are inherently linked to CI design and the number of available stimulation sites, complicating direct comparisons of spatial selectivity. Moreover, SoE values depend on the dynamic range over which responses are measured; differences in dynamic range between opsins and electrical stimulation may therefore influence the apparent sharpness of excitation profiles.

Another important limitation lies in the stimulus design. We employed relatively high optical intensities, in some cases up to 32 µJ (mW), which may lead to partial saturation of neuronal responses and thus underestimate differences in sensitivity or selectivity between opsins. At the same time, equivalent energy levels could not be applied across all opsins (e.g., ChReef), due to their lower activation thresholds, making direct comparisons less straightforward. The use of single pulses, while experimentally tractable, represents a highly artificial stimulus that does not reflect natural acoustic conditions. In addition, the stimulation durations used for optogenetic and electrical activation differ from the 100 ms pure tone stimuli used for acoustic comparisons, which we tried to compensate for by using only the onset time window in the threshold and SoE analysis but the difference in stimulus length may still affect both spatial and temporal response characteristics.

Despite these limitations, the present approach provides a useful framework for estimating the range of cumulative ICC activity across frequency bands that may be recruited by oCIs, which could ultimately inform expectations for pitch and loudness perception, as suggested by prior work (e.g. Auerbach *et al*, 2019; Vollmer *et al*, 2007). However, a more complete understanding will require moving beyond simplified stimulation paradigms and population-level measures. Future work should therefore incorporate optimized pulse parameters (e.g., durations identified in Fig. 5), lower and more physiologically relevant intensities, and multi-emitter implant designs spanning the cochlea. Importantly, responses to more complex and behaviorally relevant stimuli—such as amplitude-modulated signals, gap detection paradigms, or naturalistic sound envelopes—should be assessed. Ultimately, behavioral experiments probing pitch discrimination will be essential to validate whether the observed neural activation patterns translate into meaningful perceptual improvements.

## Material and Methods

### Animals

The presented data for f-Chrimson mediated optogenetic stimulation were recorded from 48 (25 male/ 23 female) Mongolian gerbils (*Meriones unguiculatus)* aged between 52 and 162 days. Animals were bred and housed in the animal facility at the University Medical Center Göttingen with *ad libitum* access to food and water on a 12/12 h light/dark cycle. Virus injections were performed five to nine days (6.6 ± 0.6) after birth with an Adeno-Associated Virus (AAV PHP.B-hSyn-f-Chrimson-EYFP) into the left cochlea for expressing f-Chrimson in SGNs. Electrophysiological recordings were conducted 93.4 d (± 30.7 d) post injection and were followed by euthanasia and post-mortem immunohistochemical staining and imaging for evaluation of transduction efficiency. All experimental procedures were conducted in accordance to the relevant EU regulations as well as the German Animal Welfare Act (license number: 21/00211).

### Datasets for ChReef, CatCh, eCI and acoustic click stimulation

The data presented here for the ChR variants ChReef and CatCh are taken from previous recordings (Dieter *et al*, 2019; Michael *et al*, 2023; Keppeler *et al*, 2020; Alekseev *et al*, 2025) as well as from the animals presented in the companion study (Albrecht et al). All data was reanalyzed from the raw voltage traces using the same scripts as for the newly acquired f-Chrimson data.

### Anesthesia and analgesia protocol during surgeries

All surgical procedures were performed under anesthesia using isoflurane (3-5 % for anesthesia induction, 0.5-3 % afterwards, flow rate 0.4-0.6 L/min). Breathing rate and absence of the hind limb withdrawal reflex were controlled every 15 min. The temperature was kept stable via constant monitoring and the use of custom-built heating pads. For analgesia the animals received Buprenorphine (0.1 mg/kg, every 4-6 h) and Meloxicam (adult surgeries, 0.5 mg/kg, once) or Carprofen (postnatal surgeries, 5mg/kg, once) via subcutaneous injections as well as local applications of Xylocaine spray (10 mg) on the skin before incisions and Bupivacaine drops (0.25%) on the bulla and skull before opening of the bone in the adult animals. For fluid substitution the animals received injections of Sterofundin (5mL/kg, every 4 h).

### AAV injections for gene transfer

AAV production, purification and quality control were performed as previously reported in (Huet & Rankovic, 2021). After successful anesthesia young animals (P5-P9) were injected with PHP.B_hSyn_f-Chrimson-TS-EYFP-WPRE_bGH with a virus titer of 1.3*E13 GC/mL via pressure injections (∼1 µl) into the cochlea and remained in their home cages for at least 8 weeks before the final experiment was performed.

### Stimulus generation hardware

Stimuli were delivered via a Custom-made system based on NI PCI-6229 (National Instruments, USA) controlled by custom written MATLAB scripts. Open-field acoustic stimuli were delivered from 15 cm rostrally to the animals head via a ultrasonic dynamic speaker (Vifa, Avisoft Bioacoustics, Germany) with a custom-made attenuator. We calibrated sound pressure levels using a 0.25-inch microphone (46BF-1 microphone) with a measurement amplifier (12AQ power module, GRAS Sound & Vibration, Denmark or D4039 and 2610; Brüel & Kjaer GmbH, Naerum, Denmark). For optical stimulation in the f-Chrimson animals, we used optical fibers (200 µm diameter, NA: 0.39, FT200 EMT Custom, Thorlabs, & 50 µm diameter, NA: 0.22, FG050LGA Custom) coupled to orange lasers (Coherent OBIS 594 nm LS 100 mW). Before each recording session, the fiber endings were cut (if the ending was chipped) using standard cutting tools (Thorlabs S90 R and Thorlabs XL411) before calibrating the radiant output of the emitting surface of the fiber using a power meter (Solo-2; Gentec-EO; München, Germany or S140C Thorlabs, Germany).

### Analysis toolbox

For analysis of the data, we used the custom written Matlab toolbox FEATHER v.1.0 (https://github.com/elisabethkoert/FEATHER) together with custom Matlab analysis and plotting scripts provided in the supplements of this paper.

### Auditory brainstem recordings (ABR)

We subjected animals to auditory brainstem response (ABR) measurements using 0.1 ms acoustic click stimuli at intensities ranging from 20 to 90 dB SPL in 5-10 dB steps to estimate hearing thresholds. Responses were recorded with needle electrodes placed at the vertex and behind the ear, and a reference electrode on the hind limb. For optical stimulation we accessed the temporal bone via a retroauricular incision behind the left ear and removed parts of the bulla tympanica to position a laser-coupled optical fiber in the round window of the cochlea pointing towards the cochlear apex. We then delivered optical stimuli (594 nm, 1 ms, 0–35 mW, 17 Hz) to activate SGNs and the auditory pathway. Difference potentials were preamplified using a custom-built amplifier. A bandpass filter (300-3000 Hz) was applied, and the responses were averaged across 1000 stimulus presentations. Waves were detected using a semi-automatically custom-written Matlab GUI in the FEATHER toolbox. The lowest stimulus intensity that elicited a clear wave above 3*std of the voltage trace was extracted as ABR threshold.

If no waves were detectable during the optical recordings (or if the threshold was above 10 mW), we repositioned the fiber up to four times. If there was still no ABR response, the animals were euthanized without proceeding to the midbrain recordings.

### Multi-unit Activity Recording from the Inferior Colliculus

In positive animals (oABR threshold below 10 mW), we removed the ear bones, the tympanic membrane, the facial nerve, parts of the eardrum, and the vestibular organ to enable access to the full cochlea. To access the Inferior Colliculus we performed a 1 cm longitudinal midline incision on the skull and removed all tissue in the area using forceps. For a stereotactical alignment of the skull with a micromanipulator (Luigs-Neumann SM-10), we stabilized the animal’s head by gluing a metal headpost rostrally to the bregma suture with dental cement making sure that bregma and lambda were aligned horizontally and without a lateral shift. Next, we implanted a low-impedance reference electrode (<1 Ω) on the left hemisphere between skull and dura mater and performed a craniotomy on the right hemisphere (2 mm lateral, 0.5 mm caudal to lambda) with a dental drill. After removing the dura mater we inserted a linear 32-channel silicone MEA (Acute Probe A1x32-6mm-50-177-A32, 50 µm electrode spacing, 177 µm^2^ electrode surface, Neuronexus, AnnArbor, USA) into the central nucleus of the inferior colliculus (ICC) at ∼0.2 mm/min using the micromanipulator and waited ∼30 minutes for tissue stabilization before recording, preventing brain tissue drying by applying saline drops. We acquired the neural responses at 32 kHz sampling rate and preamplified as well as bandpass filtered (0.1–8 kHz) the signals using a Digital Lynx SX recording system and the Cheetah v.6 recording software (Neuralynx, USA).

We first stimulated the cochlea acoustically using 100 ms pure tones (with 5 ms on and off ramps) between 0.5 and 32 kHz (in ¼-octave steps) at sound pressure intensities between 0 and 80 dB in 10 dB steps in a pseudo-random order to verify the MEA positioning within the tonotopic axes of the ICC. Then we proceeded to stimulate the cochlea using the 200 or 50 µm optical fibers coupled to the 594 nm Obis laser that were inserted through the round window or through cochleostomies in the midturn or apex of the cochlea. The cochleostomies were created using an electric dental drill kit (FOREDOM) and etching gel (DMG, Germany).

### Multi-unit extraction from ICC recordings

When extracting MU activity we first performed a global mean subtraction across all electrodes to remove stimulation artefacts (McInturff *et al*, 2022). Further analysis was handled for each electrode individually. The raw data traces were aligned with the trigger timepoints, bandpass-filtered (600, 6000 Hz, filter-order 4). We then approximated the standard deviation of the noise as the median absolute deviation of the baseline signal (extracted from all time windows 100 to 2 ms before trigger) divided by 0.675 (as in (Dieter *et al*, 2019; Michael *et al*, 2023; Sabesan *et al*, 2023)). Multi-unit spikes were then detected as crossings of a threshold calculated as the mean baseline activity plus three times the previously approximated standard deviation of the noise. We then extracted and peak-aligned the MU waveforms in a time window 0.5 to 1 ms around threshold crossing and extracted the peak time in reference to the trigger as spike time for the MU activity at this electrode. To avoid overestimating the response a dead time of 1 ms was introduced between detected spikes. For electrical stimulation any spikes detected within -0.5 to 2.5 ms around trigger were discarded to remove stimulation artefacts that remained despite the global mean subtraction.

### Discrimination index (d’) analysis to determine significant differences between populations of spike rates

Throughout the analysis, we had different instances where we needed to determine the difference between two sets of extracted spike rates (either when comparing stimulus and baseline activity or when comparing the response to two different stimulus intensities). For that, d’ analysis was performed (Macmillan & Creelman, 2004; Michael *et al*, 2023; Middlebrooks & Snyder, 2007). First, we created the receiver operating characteristic (ROC) curve for the two spike-rate distributions extracted from the N=30 repetitions of each stimulus. Then, we measured the area under the ROC curve (AUC) and used it to calculate the d’ value using the inverse error function (see eq. 1).

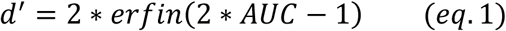

With this formula two identical spike rate distributions would yield an AUC of 0.5 and a d’=0. In the case of completely separated distributions, the AUC would get to 0 or 1 with an infinite d’. To avoid that we corrected the extreme AUC value with the ½ N procedure using eq. 2 which resulted in AUC values of 0.0003 or 0.9997 and a max d’ value of 4.88 for our recordings.

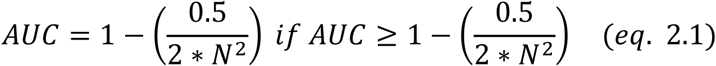

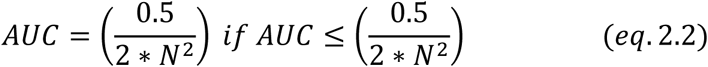

### Determination of ICC response dynamics for different stimulus modalities

We determined the ICC response dynamics for different stimulus modalities by pooling the detected spikes over multiple animals and recordings in response to high stimulus intensities for all responsive MUs. Responsive units were all those, that reached a d’ value of 1 when comparing the detected spike rates after trigger [2,25] ms with the baseline activity before trigger [-25,-2] ms for any presented stimulus intensity. We then plotted the peri-stimulus time histogram (PSTH) [–50, 150] ms around trigger for all pooled spikes within one modality. A threshold (mean + 3*std) was calculated from the baseline spike count in the PSTH at [-50,-5] ms. The response onset time t_start was set to when the first 2 bins crossed threshold, the response offset time t_stop to when 2 bins were below the threshold again with at least 2 ms between t_on and t_off to account for stimulus onset artefacts in the Multiunit extraction. The plotted PSTHs for the different modalities are shown in Fig. S4 and the extracted response time-windows are given in Table 1. For better comparison between modalities we also determined a 13 ms timewindow that only collects the strong onset-response. This starts 2 ms after trigger for optical stimuli and 5 ms after trigger for acoustic stimulation due to the faster responses observed in direct bionic SGN stimulation. For the experimental part where we investigated the response to 100 ms stimuli trains, we used a broad window encompassing the sustained response without the onset [25,125] ms.

**Table 1:** Response time windows for different stimulus modalities used in the analysis.

| Stimulus | $t_{start}$ [ms] | $t_{stop}$ [ms] |
| --- | --- | --- |
| 100 ms acoustic pure tone | 5 | 120 |
| 0.1 ms acoustic click | 5 | 25 |
| 1 ms 594 nm pulse for f-Chrimson | 2 | 18 |
| 1 ms 488 nm pulse for CatCh | 2 | 25 |
| 1 ms 522 nm pulse for ChReef | 3 | 40 |
| 0.1 ms electric biphasic pulse for eCi | 2.5 | 14 |
| 100 ms stimulus trains (optical/acoustic) | 25 | 125 |
| Onset-response optical | 2 | 15 |
| Onset-response acoustic | 5 | 18 |
| Responsive units for any stimulus | 2 | 25 |

### Determination of the ICC threshold

To determine the threshold for an ICC recording we compared the spike rates in a specific time window with the spike rates in the same time window before trigger. The lowest intensity eliciting a response that results in a d’=1 at any electrode is the threshold intensity, the corresponding electrode is the best electrode. Recordings that had an ICC activation threshold above 20 µJ for optical stimulation, above 20 mA for electric stimulation or above 60 dB SPL for acoustic stimulation were excluded, and associated with bad emitter positioning in the cochlea or suboptimal positioning of the MEA in the ICC.

### Frequency Tuning & Tonotopic Slope Assessment

After placing the MEA in the ICC we first checked the positioning of the electrodes by recording responses to pure tones and calculating the tonotopic slope. For that, we used 100 ms pure tones (with 5 ms on and off ramps) between 0.5 and 32 kHz (in ¼-octave steps) at sound pressure intensities between 0 and 80 dB in 10 dB steps in a pseudo-random order. For each frequency we determined the best electrode and ICC threshold looking at the full time window of the pure tone response [5, 120] ms. We then plotted the depth of the electrode vs the frequency and fitted a linear model to the data with two rounds of outlier removal using 1.5*std as a threshold criterion. From this fit we extracted the tonotopic slope in [oct/mm]. Recordings with a slope outside the range [3,8] oct/mm were excluded from the analysis.

### Determination of the dynamic range

We used recordings spanning a wide range of stimulus intensities to calculate the dynamic range based on the response discharge rate in the onset response time window ([2,13] ms for single pulse bionic stimulation and [5,18] ms for 100 ms pure tone acoustic stimulation).

For each stimulus and electrode, we calculated the evoked spike rate by subtracting the mean baseline spike rate (mean over the 30 stimulus presentations) from the mean spike rate in the response time window after trigger. We then plotted the max. evoked spike rate at any electrode against the applied stimulus intensity relative to the ICC activation threshold (determined using the same onset-time window). The dynamic range was extracted as the range in applied stimulus intensities in dB between reaching 10 and 90 % of the maximum of the max. evoked spike rate across all electrodes for this ICC recording.

For Fig. 2F we analysed the dynamic range for all available recordings and then filtered out to only include one recording per animal (using only 200 µm fiber RW stimulation for optical recordings) based on what recording reached the highest dynamic range for that animal. Fig. S6 shows the max. evoked spike rate for all available recordings and then gets the dynamic range based on the mean of these rates.

### Temporal Firing Behavior Analysis

To investigate the response behavior to trains of stimuli we delivered 100 ms optical trains (594 nm) through the RW at modulation frequencies from 10 to 500 Hz in 10 Hz steps using 32 mW laser power. The stimuli were constructed in a way that the first pulse occurred at 0 ms and a last stimulus occurred at 100 ms, i.e. a 50 Hz stimulus consisted of six 1 ms pulses. We determined the responsive electrodes based on the first 2-25 ms after stimulus onset and for those electrodes we extracted the spikes for the sustained response from [25, 125] ms. We calculated the vector strength (VS) for each unit by aligning all detected spikes within the stimulus cycle as previously described in (Michael *et al*, 2023) using Equation 1 with θ describing the phase of a spike and n as the total number of spikes:

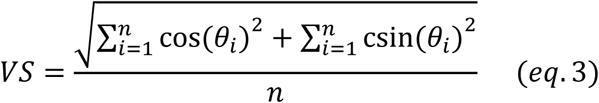

We then used the Rayleigh criterion 𝐿 = 2 ∗ 𝑛 ∗ 𝑉𝑆^2^ to set VS values that seemed to insignificant with L ≤ 13.8 (p> 0.001) to zero. For each MU we extracted the lowest rate that showed a VS of 0 and kept the previous presented stimulation rate as the maximum frequency this unit was able to follow.

In the figures we compared our results to a reference dataset from optical CatCh and acoustic click experiments on age- and sex-matched gerbils published in (Michael *et al*, 2023). For the ChReef comparison we used data collected within the companion study (Albrecht et at. 2026) and (CITE FAHDELS THESIS). All recordings were analyzed with the same analysis scripts. We only included stimulation through the round window in the optical datasets. If more than one recording existed for an animal, we automatically choose the one with most presented rates. Rates that had less than 3 recordings were removed from the dataset.

### Determination of the spread of excitation

To compare the area of activation in the ICC for different modalities we analysed 1 ms light pulse stimulation responses from our f-Chrimson recordings and from available CatCh datasets (Dieter *et al*, 2019), single electrode 0.1 ms biphasic pulses from the available eCI datasets (Dieter *et al*, 2019; Keppeler *et al*, 2020) and the pure tone recordings from the our data and the old animals. The evoked spike rate for each electrode and stimulus intensity is calculated in the onset time window (see Table 1) by subtracting the mean baseline spike rate from the mean spike rate after trigger and used to draw a evoked spike rate heatmap. The d’=1 threshold iso-contourline is added by comparing the spike rate distributions collected in the 30 stimulus repetitions from the baseline to the spike rate distribution form the response time window. The SoE is measured in the heatmap at the stimulus intensity when a specific evoked spike rate level is first reached by measuring the distance between the threshold iso-contourline at this stimulus intensity. The distance in elecrodes is then multiplied with the electrode pitch of 50 µm to get to the SoE in mm. For a visualization of the method see Fig. 4A.

### Immunohistology of Cochleae

After the recordings, animals were euthanized in accordance with animal welfare regulations. Both cochleae were removed and fixed for 1 h in 3.7% formaldehyde in PBS. Afterwards the samples were transferred into PBS and stored at 4°C.

For mid-modiolar cryostat slicing, cochleae were decalcified in 10% EDTA in PBS (pH 8.0) for up to 11 days and afterwards dehydrated using 10% sucrose for at least 24 h. Then, the samples were embedded in cryogel and sectioned into 16 µm slices using a cryostat. Around 40 sections per cochlea were collected and stored at -80 °C.

From those, selected sections were thawed, washed three times in PBS for 5 mins each, incubated in blocking buffer (GDSB, Goat Serum Dilution Buffer) for 1 h and stained overnight at 4° C with primary antibody (guinea pig anti-parvalbumin (195004, Synaptic Systems, 1:300 in 1 ml GSDB)). After primary staining, sections were washed three times with washing buffer (20mM phosphate buffer, 10% Triton X-100, 4M NaCl) for 5 mins each at RT before performing the consequent steps on injected cochleae in the dark. For secondary staining samples were incubated in AlexaFluor-labelled secondary antibodies solution (Chicken anti-GFP Alexa Fluor 488 (A-21311, ThermoFisher, 1:500 in 1 ml GSDB), goat anti-guinneapig 568 (A-11057, Invitrogen 1:200 in 1 ml GDSB)) for 1h at room temperature and washed three times in washing buffer for 5 mins each. Sections were then washed 5 minutes one time in permeabilization buffer (5mM phosphate buffer) and mounted with cover slips in Mowiol 4-88.

Cross-modiolar sections were obtained using a vibratome as described in (Thirumalai *et al*, 2025). Cochleae were decalcified in 10% EDTA in PBS (pH 8.0) for up to 11 days, prepped in 1xPBS to remove thick bone layers and embedded in 2 % agarose (dissolved in PBS) with the apex pointing upward. The samples were then cut using a vibratome (Leica 1200S, cutting speed of 0.02-0.14 mm/s, amplitude of 0.6 mm, slice thickness of 220 µm) perpendicular to the modiolus. Slices were kept in PBS until staining (0 up to 23 days). For each cochlea, 3 samples were selected representing the apex, mid and base region. Samples were blocked with goat serum dilution buffer (16% goat serum, 450 mM NaCl, 0.6% Triton X_100, 20 mM phosphate buffer, pH 7.4) for 2 h at room temperature, then incubated in primary antibodies in goat serum dilution buffer (chicken anti-GFP, 1:500, #ab13970, Abcam, USA, guinea pig anti-parvalbumin polyclonal (Cat No. 195004) 1:200 or monoclonal (Cat No. 195008), Synaptic Systems, Germany) at 4°C for 3 days. Next, samples were washed in wash buffer (20 mM phosphate buffer, 0.3% Triton X-100, 0.45 M NaCl) 3x for 10 min and incubated in the secondary antibodies in goat serum dilution buffer (goat anti-chicken 488, 1:200, A11039, Life Technologies-Invitrogen and goat anti-guinea-pig 568, 1:200, A11075, Life Technologies-Invitrogen) for 2 to 3 days at 4°C in the dark. Lastly the samples were washed 2x for 10 min in wash buffer, then 1x 10 min in 5 mM phosphate buffer, then transferred for 30 min into FOCM (30 % urea, 20 % d-sorbitol, 5% glycerol, dissolved in DMSO). The samples were mounted in FOCM on microscope slides.

All samples were imaged with a Leica SP8 confocal microscope (Leica microsystems, Germany) with a 40 x oil objective. Stacks of around 40 µm thickness with a z-step size of 1 µm and a x-y resolution of 1024x1024 pixel were acquired using the LAS-X software.

The acquired images were analysed using Arivis 4D (Zeiss) making use of custom written python scripts similar to (Thirumalai *et al*, 2025). We used the self-trained CellPose model to detect SGNs in the parvalbumin channel and applied filters for the cellsize and removed objects on the image edges. Then we placed a convex hull around all objects to get the volume of the cells. For vibratome slices this was used to calculate the 3D density values shown in Fig. S3. Since the thin cryostat slices did not allow for a good 3D analysis, we also extracted 2D density values by extracting the number of cells and area of the convex hull from a single slice in the middle of the imaged stack to calculate the values given in. For GFP transduction we extracted the intensity of the GFP channel within all SGN masks and with user input determined a threshold to separate into GFP+/-cells.

### In-silico Modeling

To obtain the example 3D structures for fiber implanted gerbil cochlea, we fixed the glass fiber positions within the cochlea using UV glue (ORBI-FlowX) and dental acrylic (Paladur) during post-mortem sham insertion surgeries. Cochleae were extracted and fixed in 3.7% FA and then transferred into PBS for imaging using X-Ray tomography with the Easytom, a commercial micro-CT system (RX Solutions, France) as described in (Schaeper *et al*, 2023, 2025). Afterwards the important cochlea structures were segmented semi-automatically using the software Amira (Thermo Fischer Scientific) as previously used in (Keppeler *et al*, 2021; Khurana *et al*, 2022). We then exported the important geometries into TracePro 7.8.1 (Lambda Research Corporation) for Monte-Carlo ray tracing. The material properties for Scala, Bone and nervous tissue (RC and modiolus) were taken from (Khurana *et al*, 2022). The fibers were modeled as glass waveguides (material properties as Fused Silica from TracePro Glass catalog). The light source was modeled as a surface source at the end of the glass fiber with Gaussian emission spectrum at a center wavelength of 488 or 594 nm (corresponding to the wavelength used in the in-vivo experiments) and a half width of 0.825 nm and an emission of 5 mW. The beam shape was modeled as gaussian pattern with x and y-half width of 20.83 or 6.5 to represent an NA of 0.5 or 0.17, respectively. For each model we simulated 10^6 rays.

As a read-out, we collected the irradiance across the RC by placing 100 small spheres of 0.01 mm along its centerline as exit-surfaces as query points (QPs). We set a threshold of 10 mW/mm2 at any point along the RC as a threshold for a targeted fiber placement and calculated the relative SoE by dividing the area under the curve for the whole RC (100 QPs long) by the peak irradiance at the best QP.

## Acknowledgements

We thank Gerhard Hoch, Christiane Senger-Freitag, Ina Preuss, Sarah Schlagowsky and Daniela Gerke for expert technical support of the study and Patricia Räke-Kügler for excellent administrative support. We thank Dres. Ursula Fünfschilling, Ramona Brecht and Jana Sie as well as Tabea Stapel for support related to preparing, executing and documenting animal experiments. We thank Dr. Lennart Roos (images of mid-modiolar sections of non-injected control cochleae), Dr. Sabina Nowakowska (images of cross-modiolar sections of non-injected control cochleae), Dr. Jannis Schaeper (X-ray tomography support), Dr. Victoria Hunniford (experimental support) and Dr. Kathrin Kusch (AAV vectors).

This work was supported and funded by Deutsche Forschungsgemeinschaft (DFG) through the Cluster of Excellence (EXC2067) Multiscale Bioimaging EXC 2067/1-390729940 (MBExC, T.Mo.) and the Collaborative Research Center 1690 (T.Mo.). The work was further supported by the Else Kröner-Fresenius Foundation via the Else Kröner-Fresenius Center for Optogenetic Therapies (EKFZ, T.Mo., B.W.) and the European Innovation Council grant OptoWavePro (101158920, TMo). E.K. is recipient of a scholarship of the Evangelisches Studienwerk e. V. Villigst. J.G. and N.A. are recipients of a scholarship of the German Academic Scholarship Foundation (Studienstiftung des Deutschen Volkes). A.V. was supported by a HORIZON TMA MSCA Postdoctoral Fellowship (OPTOCODE, grant 101107675). N.A. is recipient of a scholarship of the Goettingen Promotionskolleg für Medizinstudierende, funded by the Jacob-Henle-Programm or Else Kröner-Fresenius Foundation (2021_EKPK.04). E.K., J.G, A.V., and N.A. are collegians of the Hertha Sponer College of MBExC and A.V. and N.A. are members of the EKFZ academy.

## Supplementary material

**Supplementary Figure 1.**
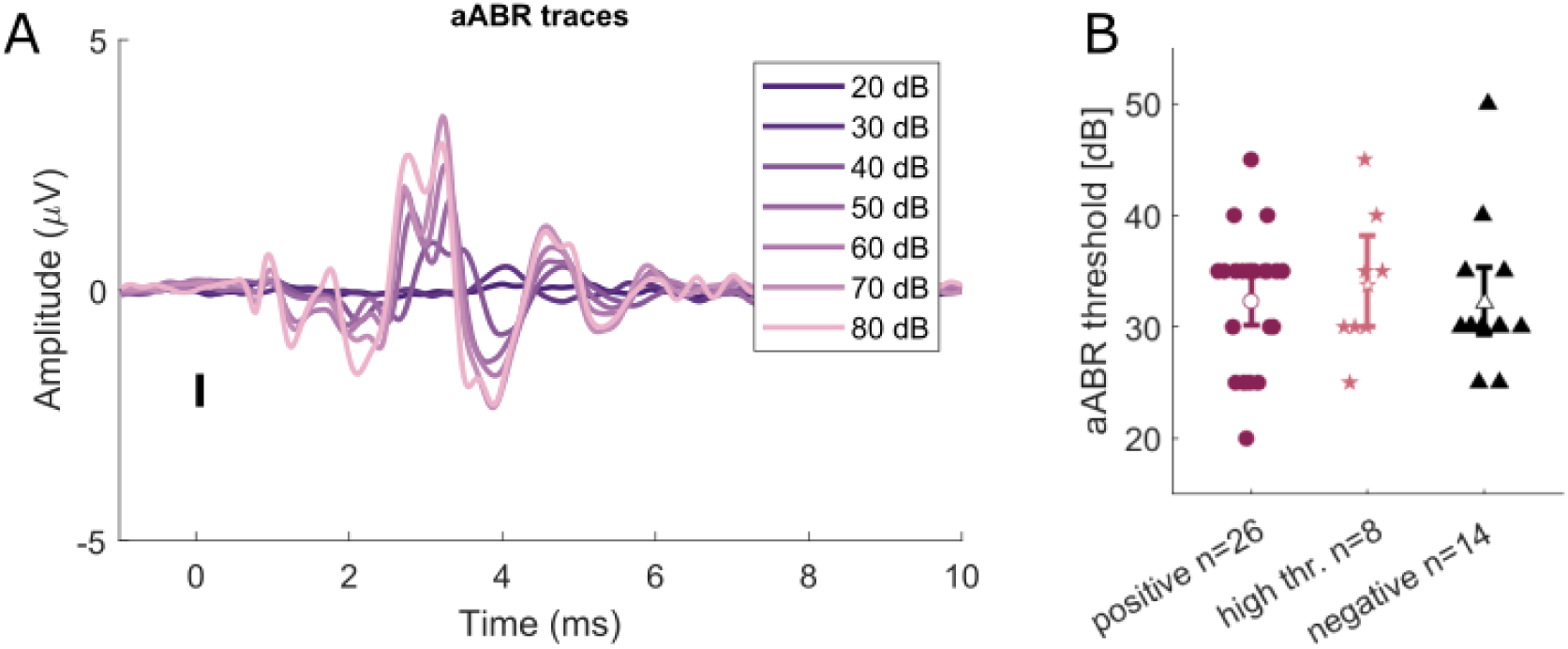
Auditory Brainstem Responses to acoustic stimulation (aABR) is not affected by the gene therapy: **(A)** Example recording of an aABR to 0.1 ms click stimuli. The thresholds was manually determined as the lowest intensity that elicited a visible wave. **(B)** aABR thresholds after separating the animals by their oABR threshodls (positive oABR <= 10 mW, high thr. oABR >10 mW, negative no response in oABR measurements for any intensity). There is no significant difference between the groups (mean ± std for positive: 32.3 ± 5.9 dB, high thr: 33.8 ± 6.4 dB, negative 32.1 ± 6.4 dB). The plot shows the mean and 95% confidence interval, for statistics a Kruskal-Wallis test with multiple comparisons was performed (*** p<0.001, **p<0.01, *p<0.1).

**Supplementary Figure 2.**
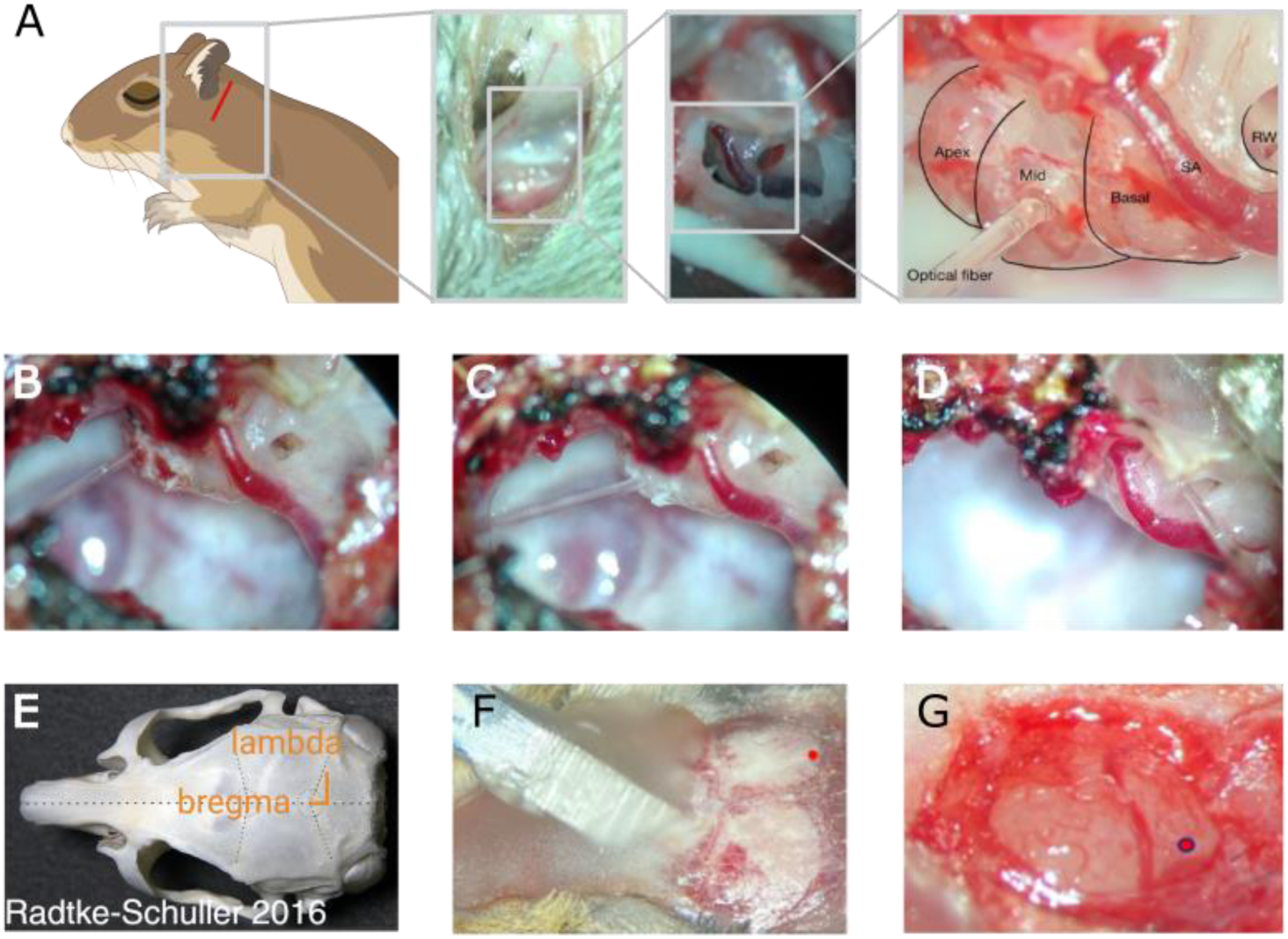
Surgical approach for cochlea and ICC access. **(A)** To access the cochlea, we performed an approx. 1cm long retroauricular incision on the left side of Mongolian gerbils under general anesthesia. After removing the skin and subcutaneous tissue, the Bulla tympanica is exposed and a bullostomy is performed exposing the cochlea with its apical, medial and basal turn as well as the round window niche (RW) and the stapedial artery (SA). **(B-D)** Fibers were placed in cochleostomies in the apex, the midturn or directly into the round window at the base to stimulate the different tonotopic areas of the cochlea. **(E)** Gerbil skull modified from (Radtke-Schuller et al, 2016) with landmarks bregma and lambda **(F)** Gerbil skull with dental cement fixated headpost as well as marking for planned drilling. **(G)** View on the brain after craniotomy with Sinus transversus at the low right end and planned marker for electrode position. (Figure created in BioRender)

**Supplementary Figure 3.**
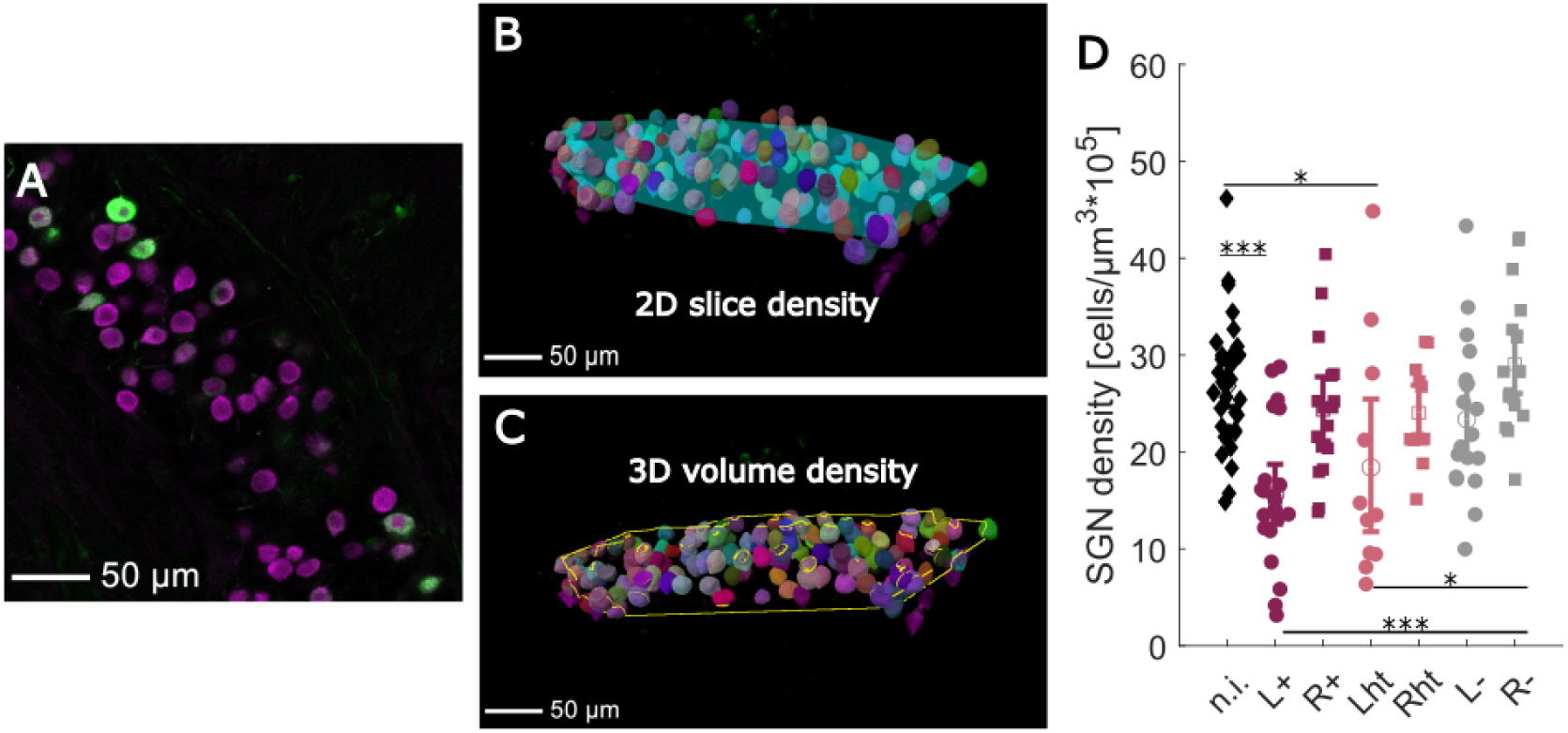
Histology analysis for 220 µm thick cross-modiolar vibratome slices allow for a 3D density estimate: **(A)** Example image taken with the 40x objective from of a cross-modiolar vibratome slice of the baseturn from an f-Chrimson injected cochlea. **(B)** Visualization of the cells detected by the Cellpose model and the intersecting area that covers all cells in the middle z-slice that was used to determine the 2D cell density **(C)** Since the imaged Rosenthal’s canal was around 40 µm thick for the cross-modiolar samples, we could also calculate the density using the full volume covering all segmented cells. **(D)** Calculated SGN density in 3D for all cross-modiolar images. The data is separated into the injected left (L) and non-injected right (R) cochlea and into animals with a positive oABR response (+) a high threshold (ht) and no oABR response (-). The non injected (n.i.) control cochleae were taken from wildtype gerbils that did not receive any gene therapy. Per cochlea we have up to 3 data points for the apex, mid and base turn. All plots show the mean and 95% confidence interval and for statistics a Kruskal-Wallis test with multiple comparisons was performed (*** p<0.001, **p<0.01, *p<0.1).

**Supplementary Figure 4.**
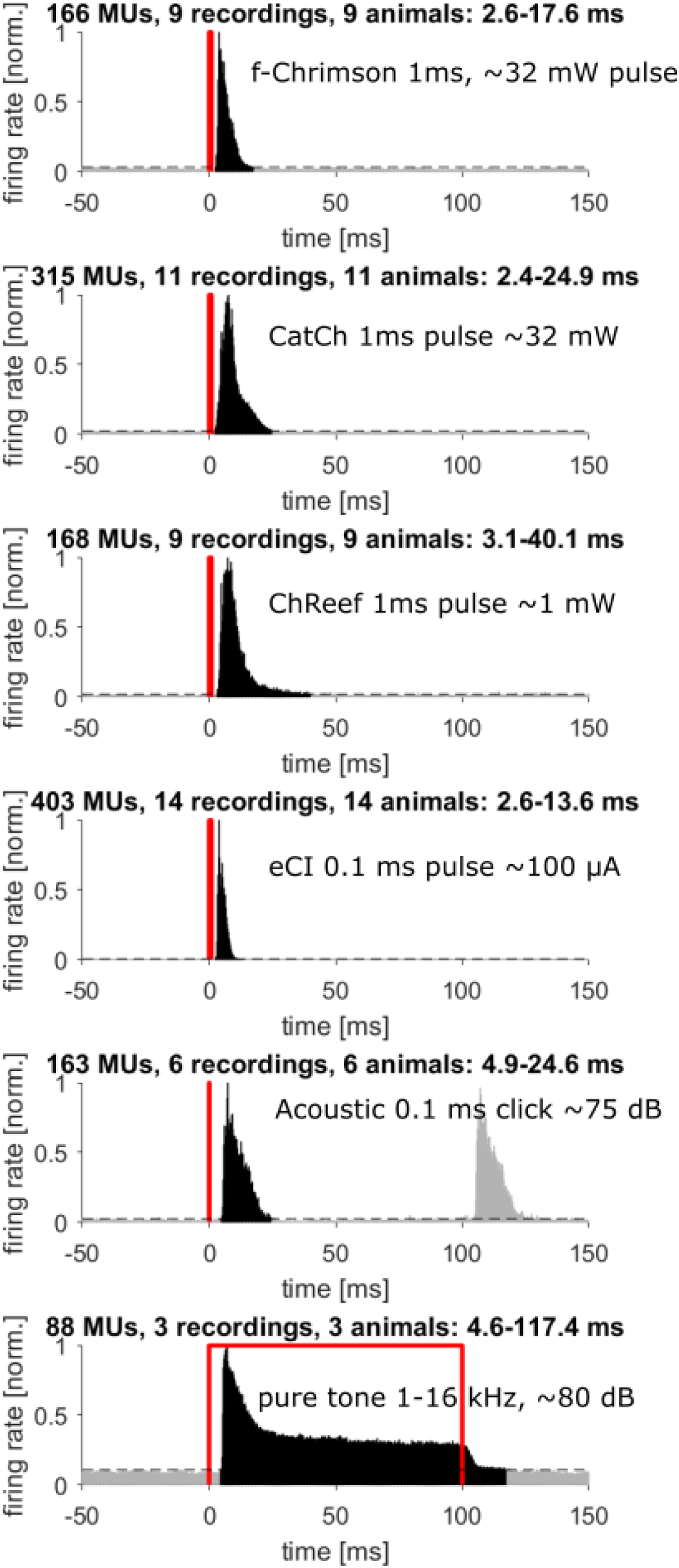
PSTHs used to determine the on/off times for each stimulus type: For the eCI data all spikes between -0.5 and 2.5 ms were removed during spike extraction to exclude the stimulation artefact. For click stimuli we did not have single click recordings so 100 ms click trains of 10 Hz were used to determine the response time window since the responses do not overlap for this low stimulation rate.

**Supplementary Figure 5.**
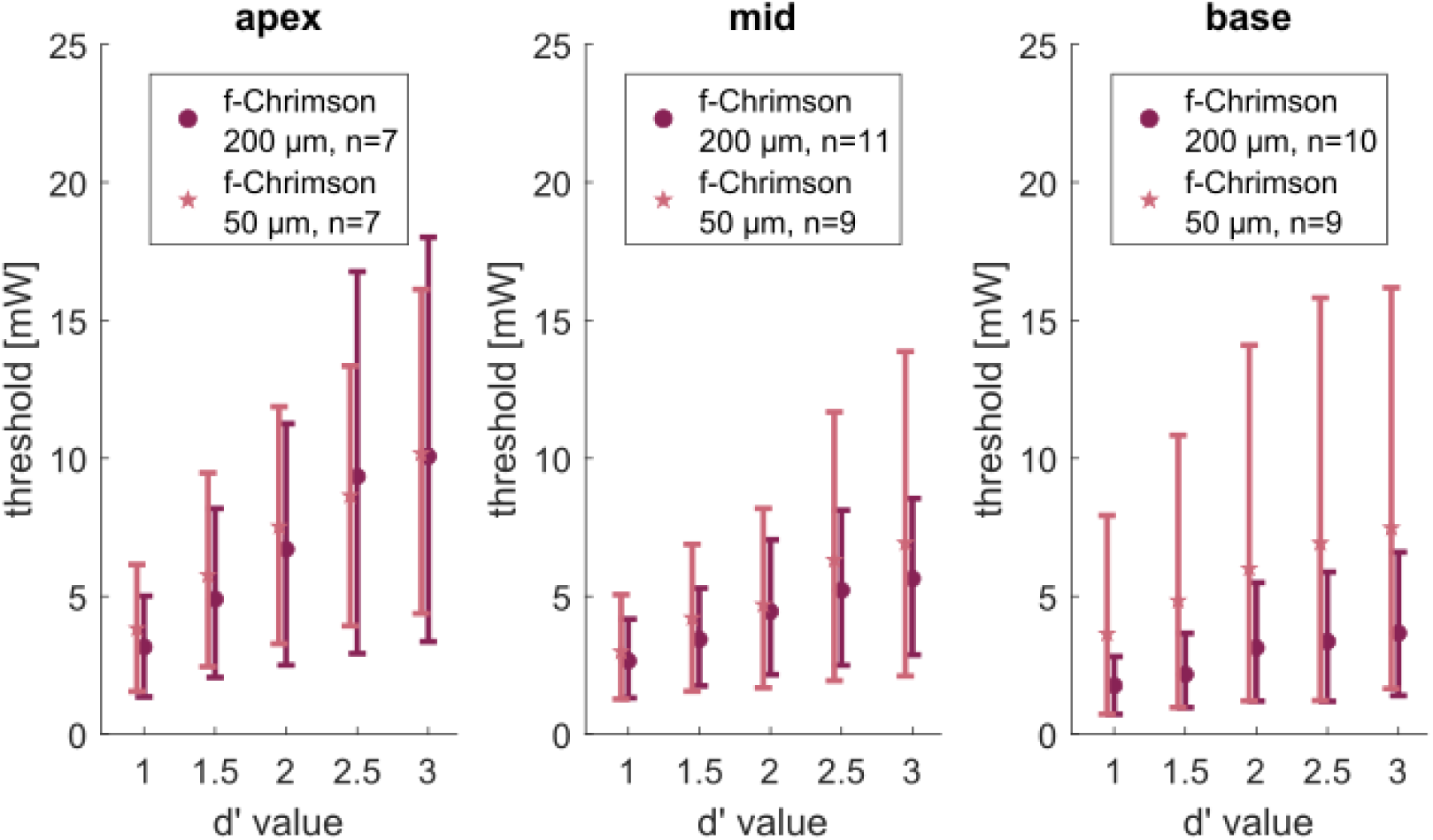
The threshold is not significantly different for different fiber diameters or cochlea positions: We performed the recordings both with 200 µm fibers and with 50 µm fibers through the same cochleostomies with a fixed fiber holder position to try to illuminate the same areas. Both fibers were calibrated on their total output in mW using an integrating sphere. We determined the threshold as the minimum intensities needed to reach a specific d’ levels when comparing the spike rates in the onset time window (2-15 ms) with the same time window before trigger. Only recordings that reached a d’ level of 3 were included and if multiple recordings existed for an animal, fiber position and diameter the one with the lowest threshold for d’=1 was chosen. Wilcoxon Ranksum testing revealed no significant differences between fiber diameters within one turn or for one diameter when comparing across turns at any of the d’ levels.

**Supplementary Figure 6.**
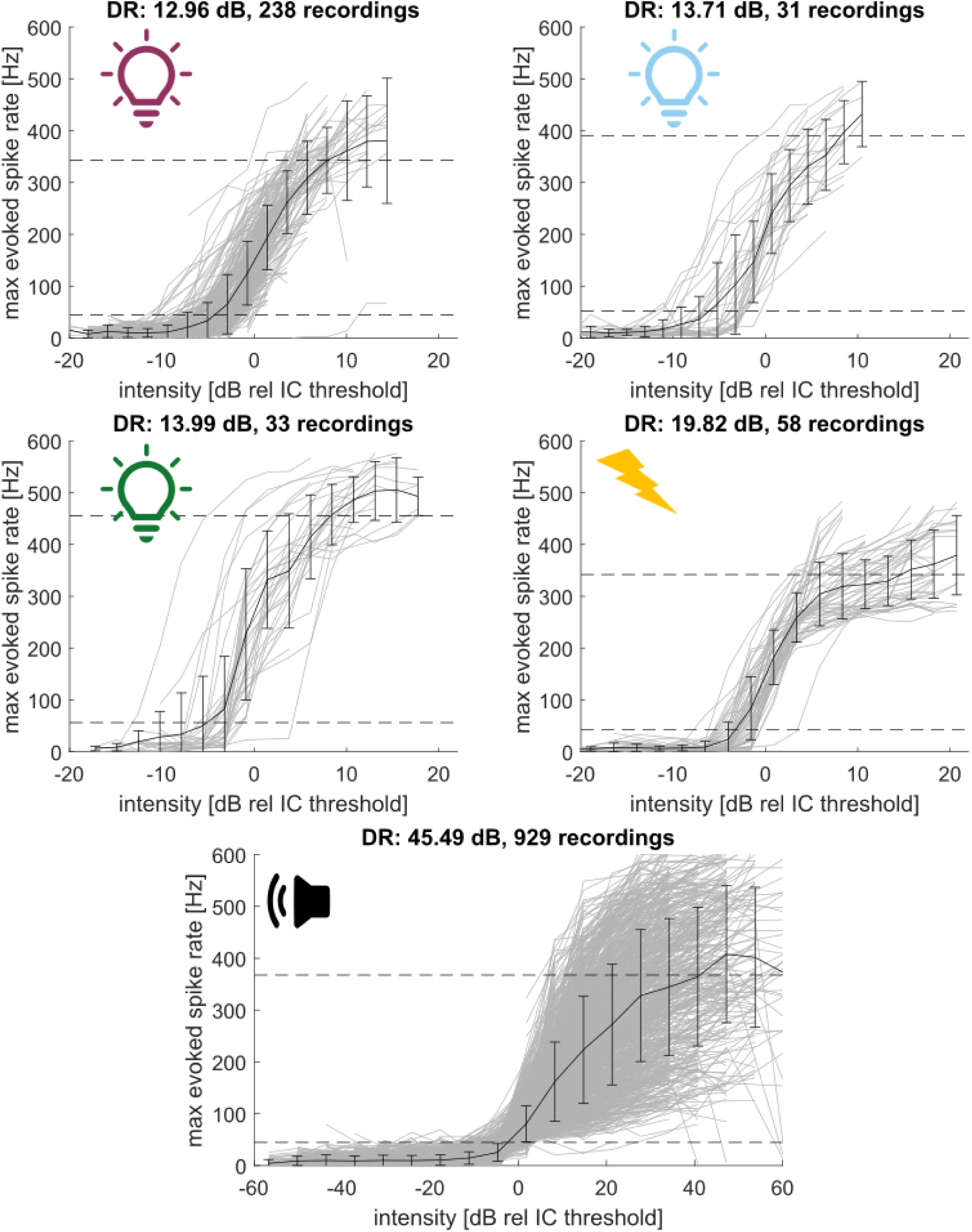
Evoked spike rates over all recordings for f-Chrimson, CatCh, ChReef, eCI and pure tone stimuli. Grey lines show the max evoked spike rate measured during individual recording at any given electrode in the onset time window (2-15 ms for bionic stimulation, 5-18 ms for acoustic stimulation). We filterered out recordings with a very high activation threshold (> 20 mW for optical stimulation, > 20 mA for eCI stimulation, > 60 dB SPL for acoustic stimulation). The black line shows the mean and std. From this mean we read out the dynamic range (DR) as the intensity range between 10 and 90 % of the maximum reached value for the evoked spike rate (dotted lines).

**Supplementary Figure 7.**
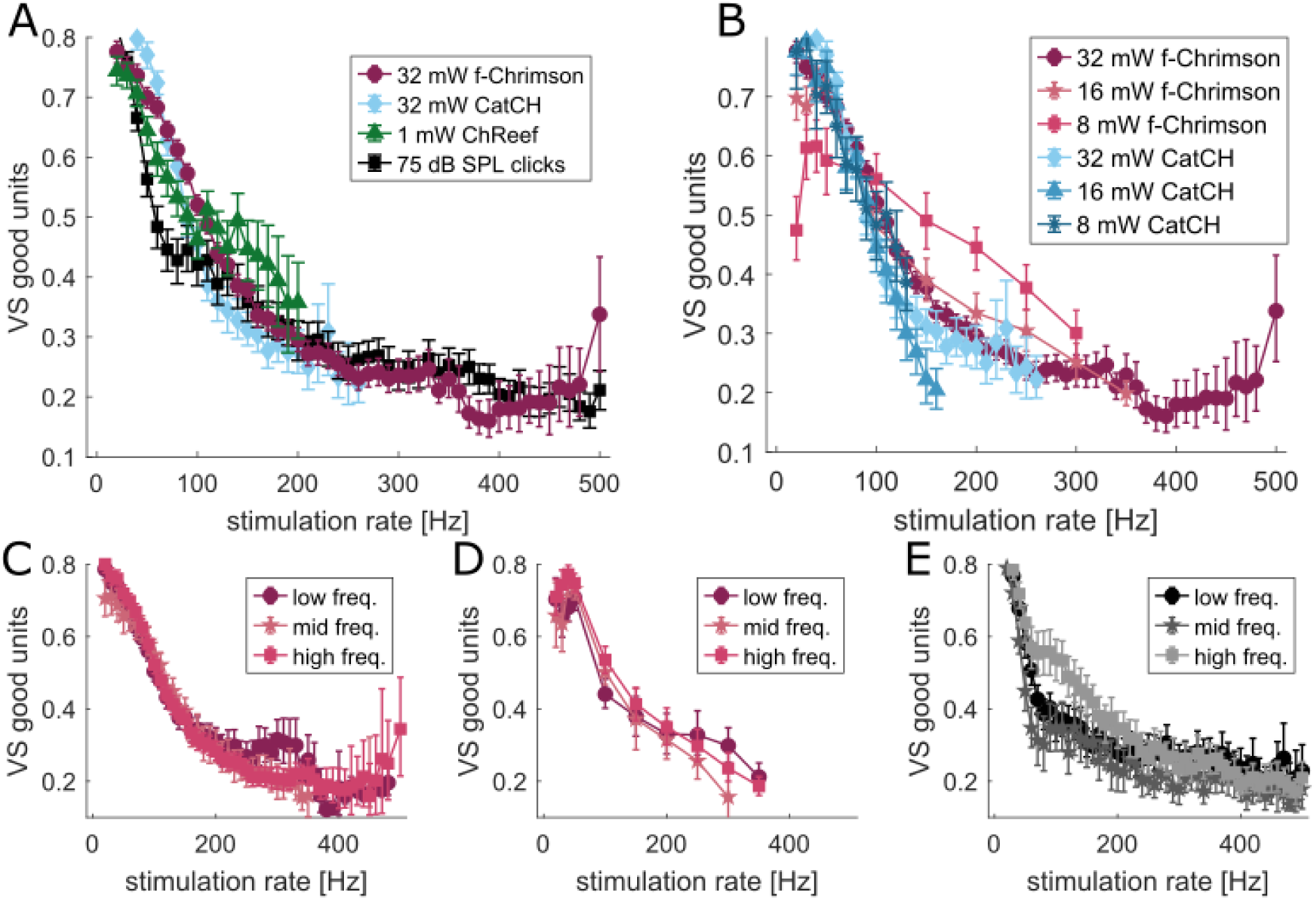
Mean VS over only those units that are able to follow a specific rate. In contrast to the VS plots given in the previous stimulation rate analysis figures, these plots only include MUs that still had a significant VS. If less then 10 MUs had enough significant VS over all animals and recordings, the mean was not plotted anymore. **(A)** Mean VS with 95% CI over a range of stimulation rate for different opsins (as in Fig. 3) only for those units that were able to follow the presented frequency. **(B)** Vector strength for only the frequency following units for the data separated by intensity plotted in Fig. EV 1 **(C-E)** VS for the frequency following units shown in Fig. EV 2 for 32 mW and 16 mW f-Chrimson stimulation as well as 75 dB click stimulation after separating the MUs according to their best frequency.

**Supplementary Figure 8.**
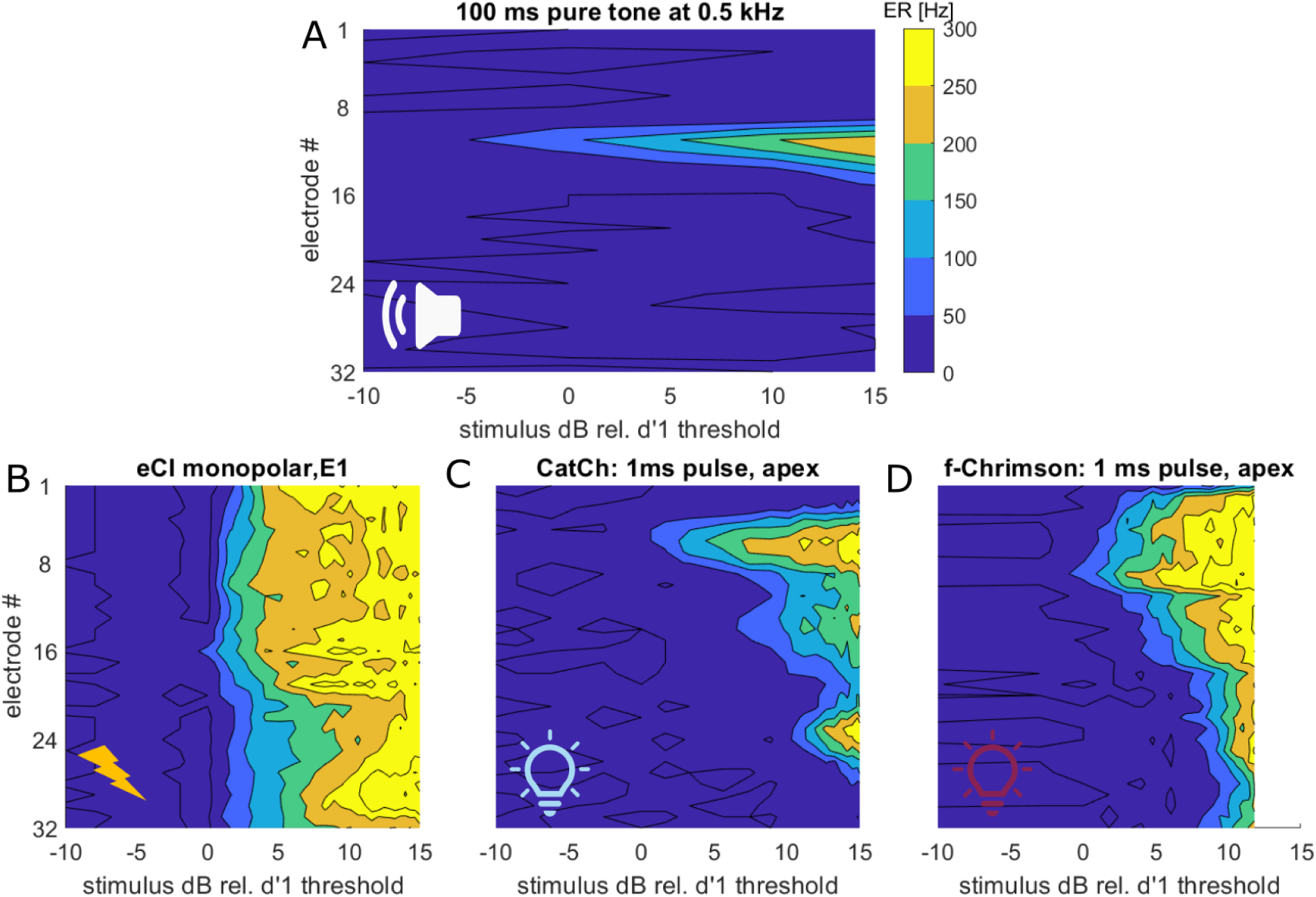
Example spike rate heatmaps for different stimulation modalities: For better visualization, we plotted the spike rate detected at each electrode within the onset time window for each stimulus modality against the applied intensity in dB rel. to the threshold intensity. As threshold intensity we used the lowest intensity at which a d’ of 1 was reached when comparing the evoked spike rates (ER) in the response time window with the baseline time window before the trigger. **(A)** response to a 500 Hz, 100 ms pure tone in an f-Chrimson injected gerbil. **(B)** response to electrode 1 in a non-injected gerbil. **(C)** response to a 1 ms, 488 nm pulse applied through an apical cochleostomy in a CatCh injected gerbil. **(D)** response to a 1 ms, 594 nm pulse applied through an apical cochleostomy in an f-Chrimson injected gerbil.

**Supplementary Figure 9.**
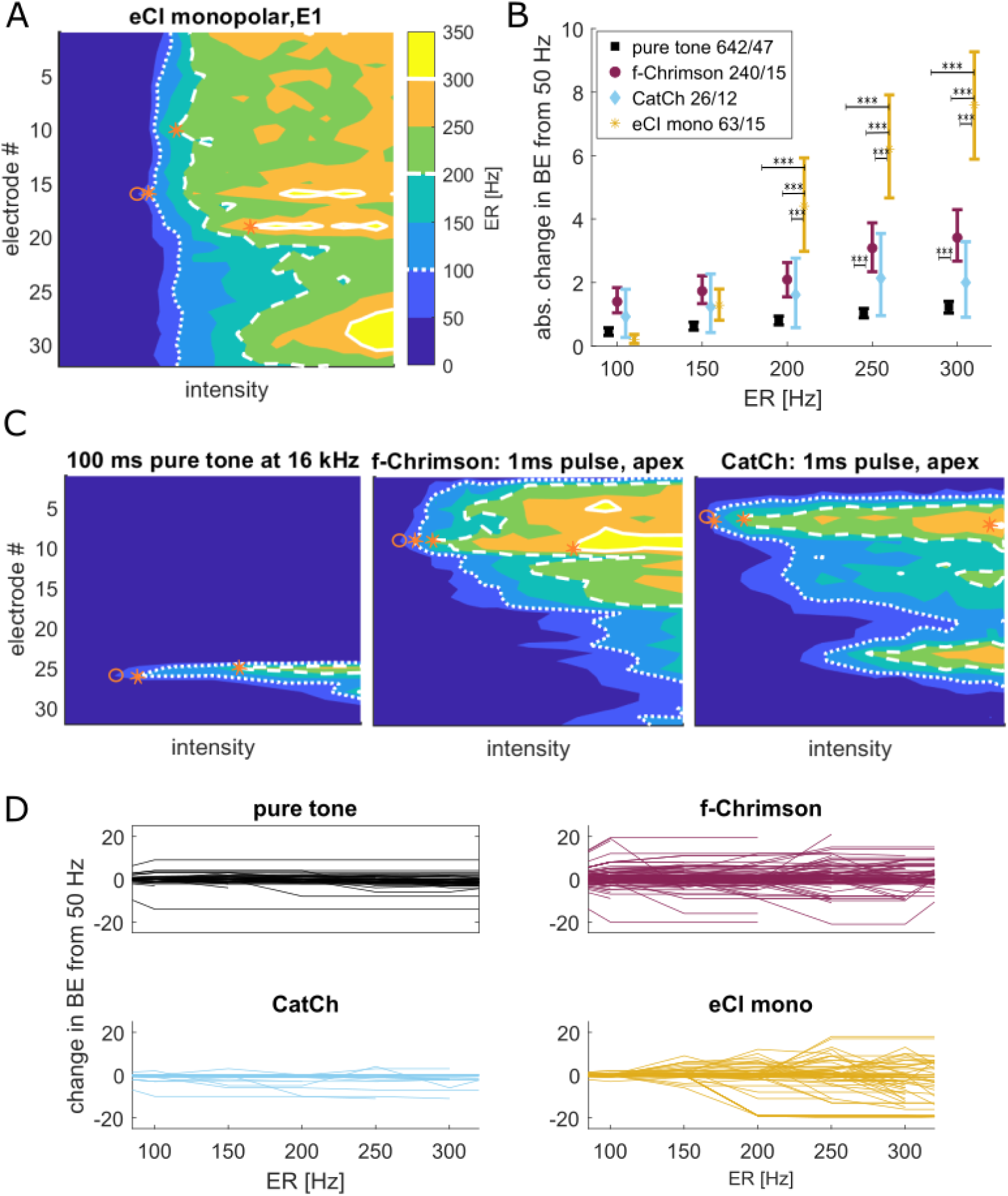
Electrical stimulation shows a stronger change in area of peak activation with higher intensities than optical or acoustic stimulation: When observing the evoked spike rate over a range of intensities, the best electrode can change position along the depth of the ICC for increasing d’ levels. **(A)** evoked spike rate (ER) values for an eCI recording, where the best electrode moves from elec. 16 at 50 and 100 Hz to 10 and to 19 for 200 and 300 Hz evoked spike rate as readout level. **(B)** absolute change in electrodes from the best electrode for 50 Hz when moving to higher evoked spike rates for all stimulus modalities. The plots show the mean and 95% CI. We performed a multiple comparison ANOVA (ranova and multompare matlab functions) to test for significance between the groups (* p<0.05, ** p<0.01, ***p<0.001). **(C)** same evoked spike rate plots as in A for a pure tone (BE starts at 26 at 50 Hz and moves to 25 for 200 Hz), f-Chrimson (BE for 50, 100 and 200 Hz at 9 for 300 Hz at 10) and CatCh (BE at 5 for 50 Hz, at 6 for 100 and 200 Hz and at 7 for 300 Hz) recording. **(D)** actual drift in best electrode for all recordings included in the analysis (for pure tones only every fifth is plotted) to show in which direction the peak of activation drifts. Positive values mean the new BE are located higher in the ICC which means a shift of stimulation towards the apex/ low frequency regions.

**Supplementary Figure 10:**
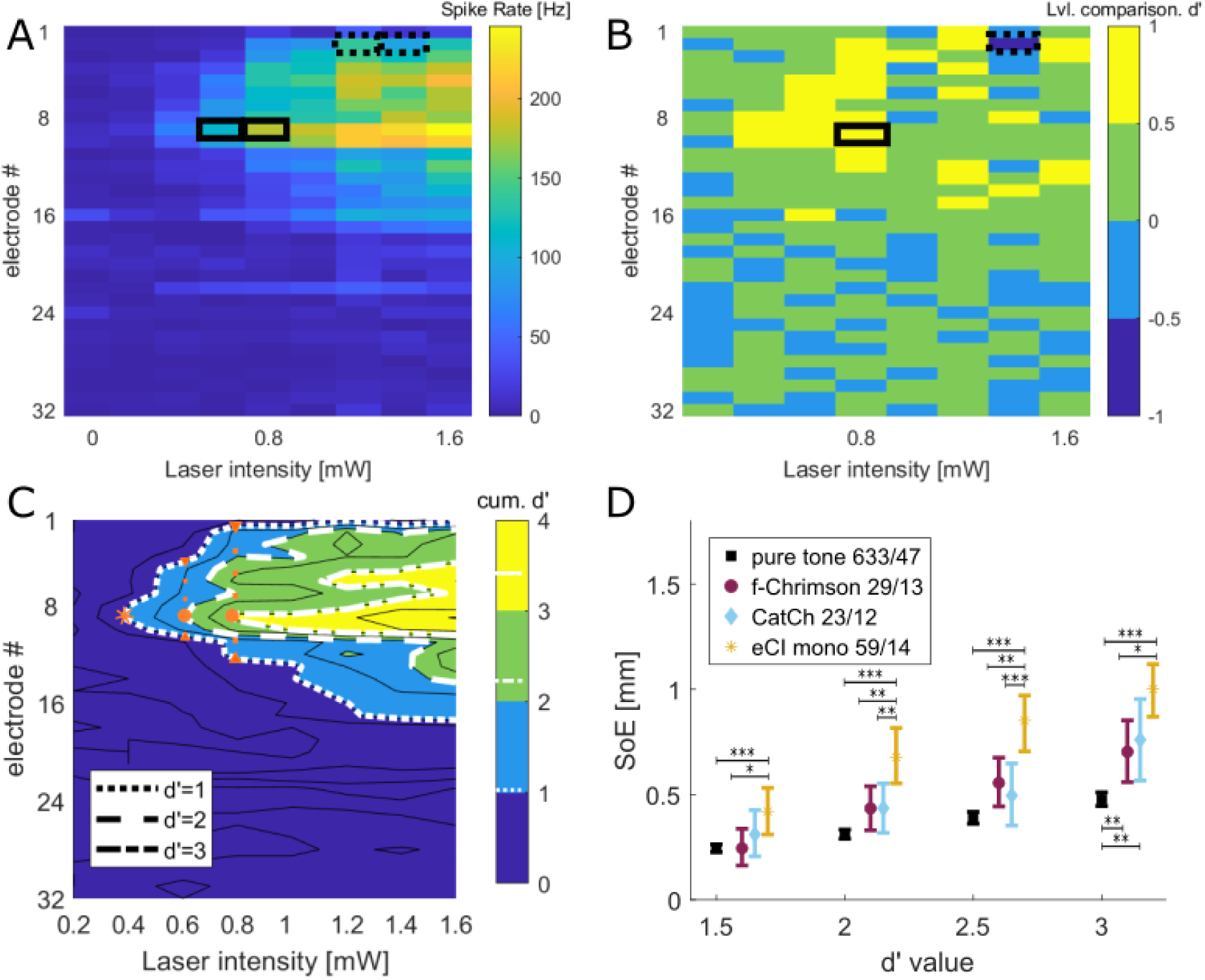
Using fully dPrime based SoE calculation as in (Dieter et al, 2019) To better align our results with previous publications from the lab, we also implemented a function that determines the SoE in a similar way. The spike rates for each electrode and intensity are calculated for all 30 stimulus repetitions in a modality specific time window based on the PSTH (as given in Table 1). **(A)** shows the mean spike rate heatmap. Then, at each electrode, we use d’ analysis to compare the spike rate distributions of the increasing levels of stimulation. **(B)** shows the d’ heatmap. When the mean spike rate increases (boxes with full lines) the comparison d’ will be positive, if the spike rate decreases for the higher stimulus intensity, the d’ value will be nagtive (dotted boxes). **(C)** We sum up the d’ values along each electrode to get a cumulative d’ value, that illustrates the change in spike rate for each electrode over the applied range of intensities. When plotting these cum. d’ values in a contour heatmap, we can measure the width of the d’=1 isocontour line at different intensity readout levels. Here the readout intensity is determined by when the best electrode (elec. 9) first reaches a specific cum. d’ value. **(D)** Mean SoE for the recordings shown in Fig. 4 when applying this type of SoE calculation. The numbers in the legend show number of MUs/animals. Only recordings that reached a cum. d’ of 3 Hz were included. If multiple recordings existed for a diameter and cochlea position the one with the lowest spread at cum. d’=1.5 was chosen.

**Supplementary Figure 11.**
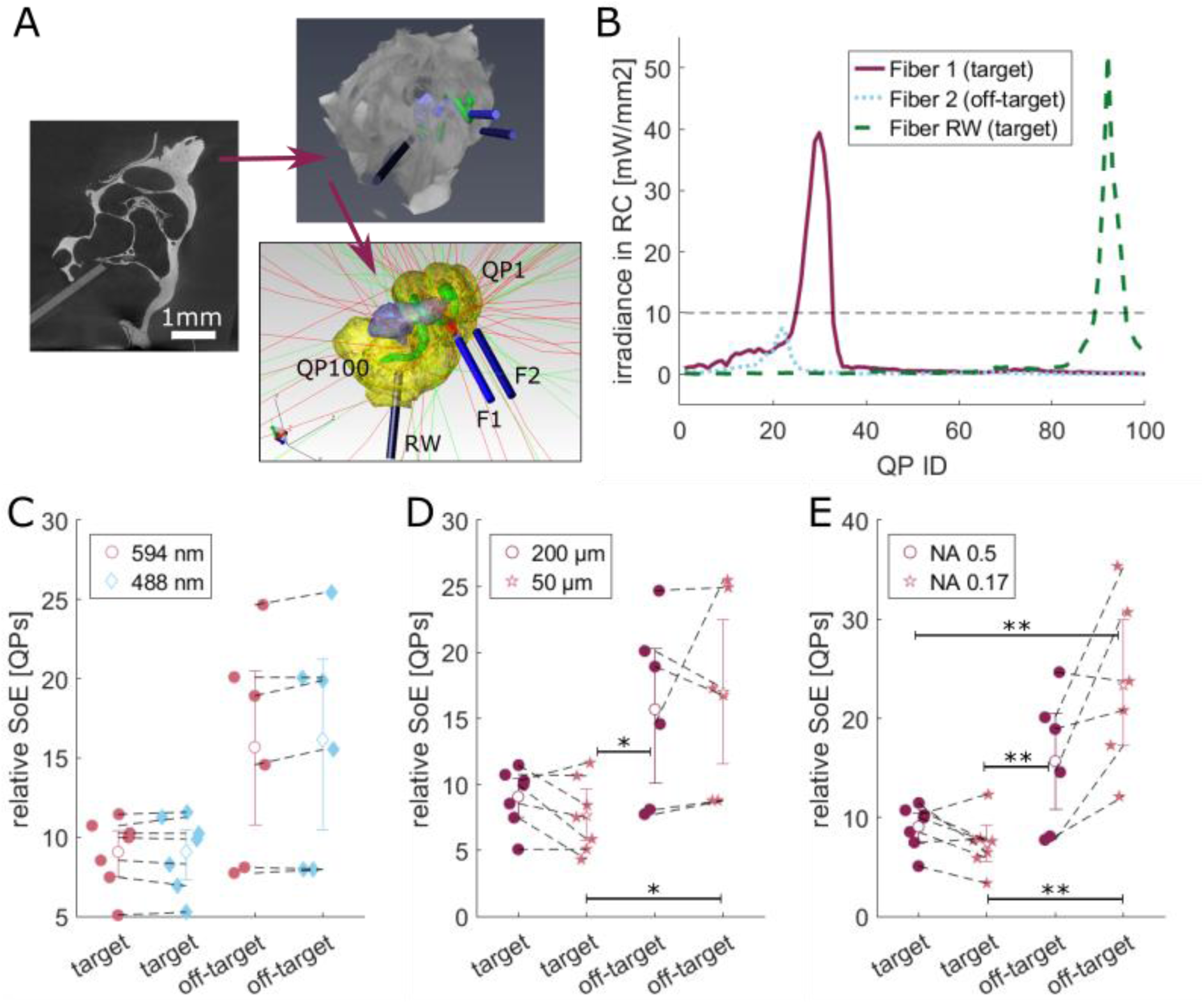
Light path simulations show that fiber positioning is more critical for the spread of light in the cochlea than the light wavelength, numerical aperture or fiber diameter: We performed sham insertions on sacrificed animals and glued the fibers into position to have examples with varying insertion parameters to optimize the fiber positioning. This sample set of n=7 cochleae with 1 to 3 fibers each was used to test the influence of fiber and light properties on the spread of light in the cochlea using computer simulations. **(A)** The recorded X-ray tomogrophies were used to reconstruct the fibers (blue) as well as central cochlea structures (scalae-yellow, Rosenthal’s canal (RC)-green, modiolus-purple) in Amira. Based on these surfaces, a TracePro model was built to simulate the irradiance of the different fibers across query points within the center of the RC with QP1 at the apex and QP100 at the base. **(B)** The measured irradiance values for the different fibers in A along the RC. A threshold of 10 mW/mm2 was chosen to separate the fiber positions into target fibers that point towards the RC (here F1 and RW) and off-target fibers that mostly miss the RC (here F2). This simulation used 594 nm light, 200 µm fibers and a numerical aperture (NA) of 0.5 (default values). For a relative measurement for the spread of excitation by the light the area under the curve (AUC) was determined and divided by the peak irradiance. These numbers are shown in **(C,D,E)** with one parameter changed from the default values. The lines connect the results for the same fiber with only the one variable change indicated in the legend. Ranksum tests were performed between all groups individually (**p<0.01, *p<0.1).

**Supplementary Figure 12.**
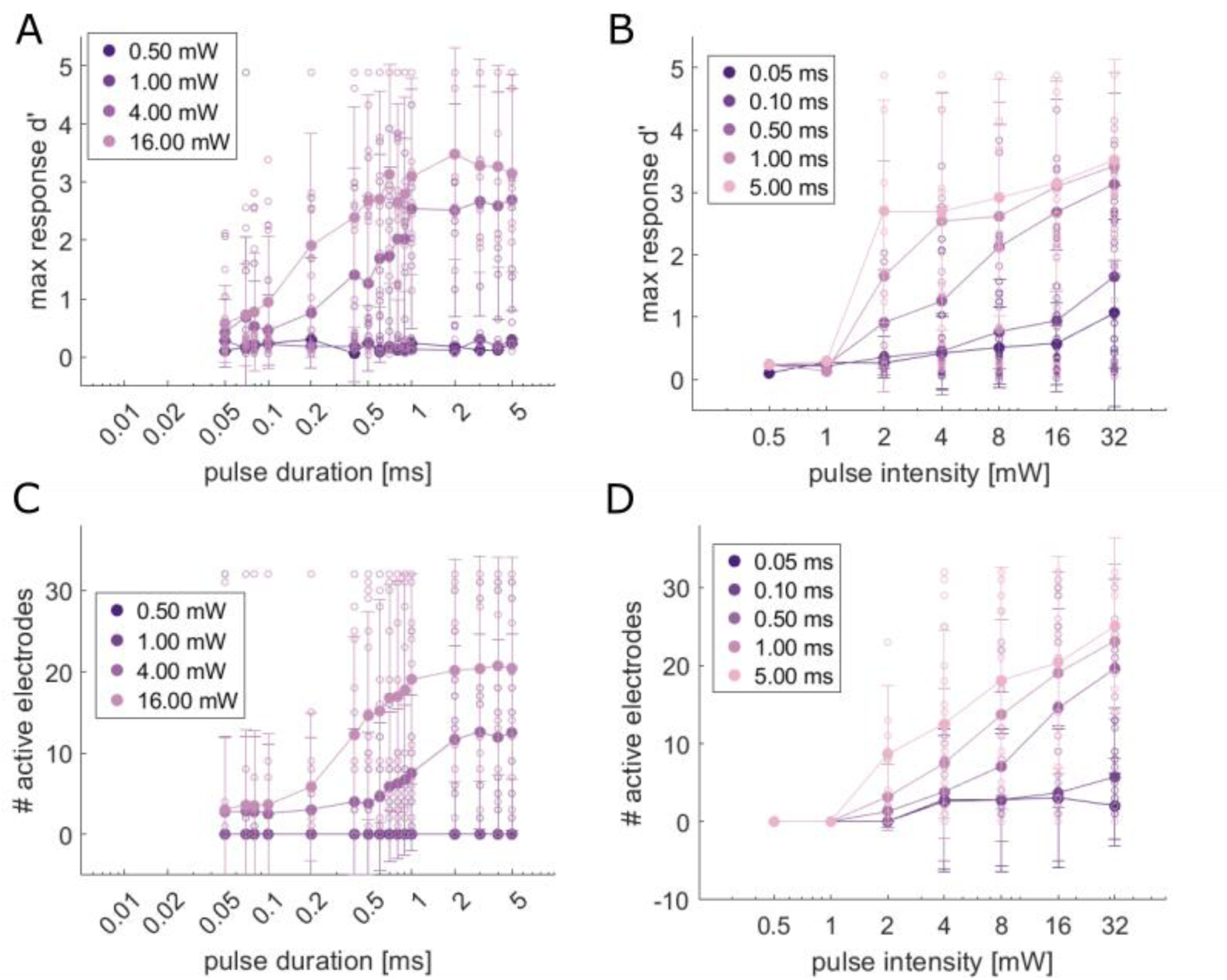
Detailed effects of pulse duration and intensity and duration opn the response strength and number of active electrodes: The plots show the individual datapoints as well as the mean and std. for the 16 animals that were presented with the pulse duration-intensity stimulation protocol. **(A)** and **(B)** show the max d’ value (value from the one best electrode) that is reached when comparing the spike rate in the response time window with the baseline rate before trigger onset. These values were used to generate the means presented in Fig. 5C. **(C)** and **(D)** show the number of active electrodes (that reached a d’ above 1 for the comparison with the baseline rate) for the different stimulus conditions.

